# Predicting protein-membrane interfaces of peripheral membrane proteins using ensemble machine learning

**DOI:** 10.1101/2021.06.28.450157

**Authors:** Alexios Chatzigoulas, Zoe Cournia

## Abstract

Abnormal protein-membrane attachment is involved in deregulated cellular pathways and in disease. Therefore, the possibility to modulate protein-membrane interactions represents a new promising therapeutic strategy for peripheral membrane proteins that have been considered so far undruggable. A major obstacle in this drug design strategy is that the membrane binding domains of peripheral membrane proteins are usually not known. The development of fast and efficient algorithms predicting the protein-membrane interface would shed light into the accessibility of membrane-protein interfaces by drug-like molecules. Herein, we describe an ensemble machine learning methodology and algorithm for predicting membrane-penetrating amino acids. We utilize available experimental data in the literature for training 21 machine learning classifiers and a voting classifier. Evaluation of the ensemble classifier accuracy produced a macro-averaged F_1_ score = 0.92 and an MCC = 0.84 for predicting correctly membrane-penetrating amino acids on unknown proteins of an independent test set. The python code for predicting protein-membrane interfaces of peripheral membrane proteins is available at https://github.com/zoecournia/DREAMM.

## Introduction

Membrane proteins are topologically divided in transmembrane proteins that are permanently attached in the interior of the membrane, peripheral membrane proteins that associate non-covalently with the surface of the membrane, and lipid-anchored proteins that attach to the membrane with a covalent bond [1]. Peripheral membrane proteins are essential in cellular processes such as transporting substances across the cell membrane, activating proteins and enzymes, regulating signal transduction, and other functions [1, 2]. Abnormal protein-membrane attachment due to membrane-binding domain mutations and peripheral membrane protein over- or under activation are involved in deregulated cellular pathways and in disease [1, 3–8]. Hence, the possibility to modulate protein-membrane interactions represents a promising therapeutic strategy for many disease indications and in particular for targeting membrane proteins that have been considered undruggable such as the membrane-anchored KRAS protein, which is implicated in over 30% of cancer types [9, 10], α-synuclein, which is a main pathological hallmark of Parkinson’s disease [11, 12], and lipid kinases such as PI3Kα, which is the most frequently mutated kinase and present in a variety of tumors [13] with one of its hotspot mutations, H1047R, acting on altering the protein’s association with the cell membrane [14–17].

The feasibility of targeting protein-membrane interfaces is supported by the fact that peripheral membrane proteins contain a membrane-binding domain with cavities that could be potentially targeted by small molecules [18, 19]. The literature reports the feasibility of targeting the protein-membrane interface indicating that therapeutic targets binding transiently to the membrane can be targeted with small molecules, and that inhibitors of protein-membrane interactions may be identified [18, 20–25]. However, these examples are only limited compared to the overall drug design efforts of the community indicating that the accessibility of protein-membrane interfaces by small molecules has been so far unexplored possibly due to the complexity of the interface, the limited protein-membrane structural information, and the absence of tools and workflows to automate the drug design process at the protein-membrane interface. Moreover, protein-membrane interaction sites of peripheral membrane proteins are commonly undiscovered; hence, the first step into modulating the protein-membrane interface is their identification.

Several efforts towards the design of tools that detect protein-membrane regions, domains, and lipid-binding sites have appeared [26–29]; however, these are mainly applied to directly to 1D protein sequences without considering the protein structural information and in many cases the web links are outdated [26–28]. To our knowledge, only two methodologies, which predict these interaction sites from the 3D protein structure, are currently publically available: the Positioning of Proteins in Membrane (PPM) [30, 31] and the Membrane Optimal Docking Area (MODA) [32]. PPM combines an anisotropic solvent representation of the lipid bilayer, an all atom representation of a solute, and a universal solvation model, calculating rotational and translational positions of transmembrane and peripheral membrane proteins in membranes [30, 31]. MODA is based on the protein-protein interface predictor PIER [33], which builds a set of evenly distributed points at 5 Å from one another and from the protein surface, defining each patch as the set of all protein surface atoms. Then, it calculates a score based on atom solvent-accessible surface area (SASA) and atom type specific weights, and transfers the patch membrane propensity scores to surface amino acids, thereby predicting which amino acids contact the cell membranes.

Herein, we present an automated prediction algorithm using ensemble machine learning, which identifies membrane-binding interfaces with high accuracy (macro-averaged F_1_ score = 0.92 and an MCC = 0.84) taking as input the 3D peripheral membrane protein coordinates and demonstrates better accuracy than existing methods.

## Methods

### Data Preparation

To construct the dataset, we use 54 peripheral membrane proteins with known 3D structures and experimentally known membrane-penetrating amino acids, retrieved from extensive literature search. For the dataset generation, protein structures were prepared by deleting unwanted chains and co-crystallized solvent atoms, adding missing side chain atoms, and converting non-standard amino acids to their standard equivalents using HTMD [34]. In case of NMR-resolved structures, the first model of the NMR ensemble was kept. Then, the dataset was split in a training set (~85% of the dataset, Table S1) and a test set (~15% of the dataset, Table S2). Finally, a dataset of 12.805 amino acids consisting of the training set samples and a dataset of 2.177 amino acids consisting of the test set were assembled. These samples were labeled in two classes, the membrane-penetrating and the non-penetrating amino acids, leading to a highly imbalanced binary classification problem (supervised learning), where the membrane-penetrating amino acids comprise ~1.3% of the total samples in the training set. The feature extraction, and feature and data selection processes are explained in detail in the SI.

The most important features were expressing hydrophobicity and solvent exposure, but also: evolutionary conservation, secondary structure (coil loop or not), flexibility (squared fluctuations calculated with the Gaussian network model, probably associated with the flexible coil loops), dihedral angles (which again might be associated with the secondary structure), and Transferable Atom Equivalent (TAE) descriptors [35] (which are molecular properties related to electron density distribution) were determined to be critical for the accuracy of the results (more information in the SI, in Figure S1, and in Table S3). To balance the two classes several techniques were utilized leading to six different balanced datasets (see SI for more information and Figure S2).

Furthermore, a second test set of 11 peripheral membrane proteins with known 3D structures and experimentally known membrane-penetrating regions (not amino acids) was assembled. As no specific amino acids were experimentally tested, these proteins were assessed qualitatively as a second test set.

Finally, to ensure that the predictions are unbiased, the percentage of identical amino acids of the pairwise sequence alignment was calculated for all sequence pairs of the dataset, revealing high percentage identity values (more than 40%) for proteins in the training set, but not in the test sets, ensuring that the predictions in the test sets are unbiased (more information in the SI and Tables S4 and S5).

### Ensemble machine learning methodology

For each one of the six training sets, 21 machine learning classifiers were trained: 19 from the scikit-learn Python package [36], the LightGBM classifier [37], and the XGBoost classifier [38]. The hyper-parameters of these classifiers were optimized to discover the best hyper-parameters that fit the data for each classifier (Table S6) [39, 40]. Specifically, for every training set and for every classifier the randomized search cross-validation technique was performed using 5 folds in a wide range of hyper-parameter values, training hundreds of thousands models (Table S6). Subsequently, iteratively exhaustive searches were performed (grid search cross-validation with 5 folds) in a small range of hyper-parameter values in the vicinity of the best hyper-parameter space determined from the randomized search cross-validation, which led to a set of optimal hyper-parameters. Notably, in the over- and under-sampled datasets, over- and under-sampling was performed in each fold separately to avoid information leak in the validation fold.

To assess the performance of the classifier models for both the randomized and grid search cross-validation procedures, the macro-averaged harmonic mean of the precision and recall, F_1_ score, was chosen as a metric. Recall expresses the amount of correct predicted true positives (Eq. 1), while precision expresses the predicted true positives that are actually true (Eq. 2). The general formula of F score is derived based on a positive real variable β, where β determines the importance of recall over precision (Eq. 3). When β = 1 (F_1_ score), recall and precision are weighted equally (Eq. 4), when β < 1 more weight is given in precision and when β > 1 recall is favored.

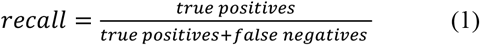

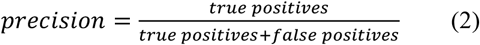

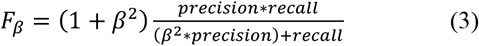

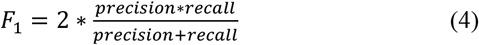

For more information about the hyper-parameter tuning process and the performance metrics please refer to the SI.

Subsequently, the resulting predictions of the aforementioned classifiers were imputed as input to a meta-classifiers (second-level classifier). The voting classifier, which classifies a sample based on the majority voting of the first-level classifiers [41], and the stacking classifier, which trains a classifier on the output of the first-level classifiers in order to compute the final prediction [42], were employed using the Python library mlxtend [43]. In both meta-classifiers, all possible combinations of the first-level classifiers were examined to discover the best classifier combination. Every classifier combination was tested using the independent test set with known protein-membrane amino acids (Table S2) to measure the combination with the best performance. Subsequently, considering that not every amino acids in the dataset was experimentally tested, resulting in membrane-penetrating amino acids marked as non-penetrating, the best models were manually inspected in order to assess their false positives, and the final model was chosen based on F_2_ score (see Results). A schematic representation of the above procedure is illustrated in Figure 1.

**Figure 1.**
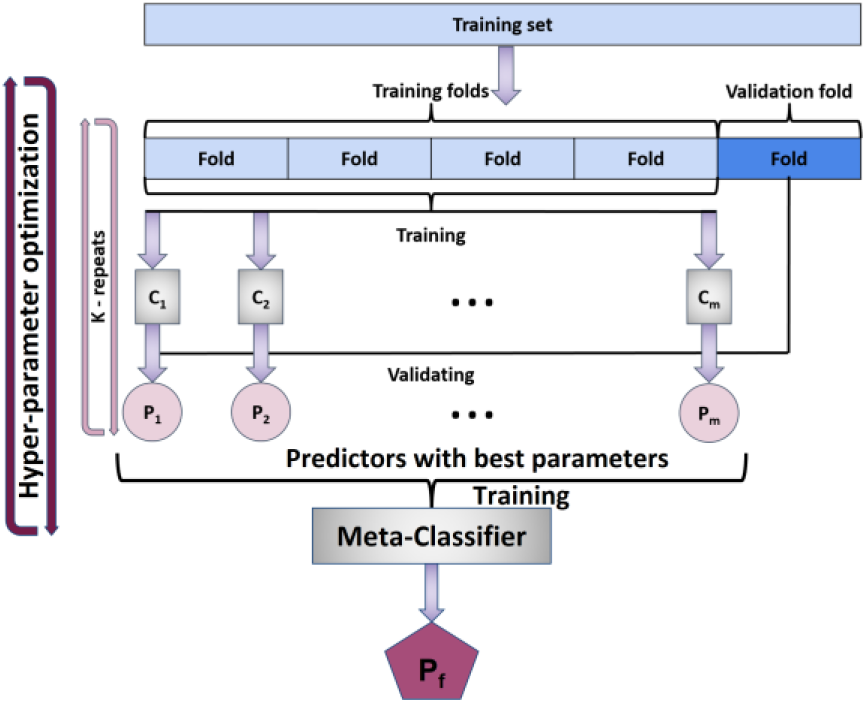
For each of the six datasets, we optimized the hyper-parameter space of 21 classifiers using 5-fold cross-validation on the training set. The predictions of the models with the best F1 score from these classifiers were provided as input to meta-classifiers. Given the F2 score on the test set the best meta-classifier was kept as the final predictor.

## Results

The first-level classifiers providing the most accurate results were those trained in the initial dataset using weights. For these classifiers and initial dataset, several second-level classifiers exhibited better performance than the individual classifiers in terms of F_1_ score, precision/recall area under the curve (PR AUC), Matthews correlation coefficient (MCC), and other metric scores. The Receiver Operating Characteristics (ROC) AUC, which is regularly used in literature, was not considered as it may be misleading for highly imbalanced classification problems such as our case (see the SI) [44, 45]. Then, results from the top second-level classifiers were subject to manual inspection and the best was selected according to the F_2_ score to emphasize on recall. Although, seemingly, it is natural to prioritize on precision as in our case false positives are more critical than false negatives, manual inspection of the false positive results of the top second-level meta-classifiers indicated that these could actually be true positive membrane-penetrating amino acids as they are adjacent to amino acids that are membrane-penetrating or aligned with them to adjacent loops (see below). Finally, the best performing second-level classifier was the voting classifier for a combination consisting of five classifiers, the linear discriminant analysis, the logistic regression, the linear support vector classifier, the decision tree classifier, and the light gradient boosting machine. Various metric scores of the 21 first-level classifiers and the chosen second-level classifier for the initial dataset using weights can be viewed in Table S7.

The test set predictions can be viewed in Figure 2 and Table S8, where ~2/3 of the false positive amino acids are in fact correct predictions as they are located in the protein-membrane interface adjacent to true positives or on adjacent loops. For example, in retinoid isomerohydrolase, amino acid F262 is in a 4 Å distance from the experimentally confirmed membrane-penetrating amino acids; for the glycolipid transfer protein, amino acids I143 and Y153 are next to and aligned with W142. In other examples, i.e. the cholesterol-regulated Start protein 4, amino acid M196 although it is located in different loop, it is aligned with L124, as well as for the phosphatidylinositol transfer protein beta isoform, where M74 is in a different loop but aligned with the experimentally-confirmed membrane-penetrating amino acids W202 and W203; for the PH domain of the ceramide transfer protein, I37 and W40 are next to W33 and Y36, and F81 is aligned with them in an adjacent loop. Considering that these predictions are in fact located at the protein-membrane interface, they can be considered as true positives and the macro-averaged F_1_ score increases from 0.86 to 0.92 and MCC from 0.71 to 0.84.

**Figure 2.**
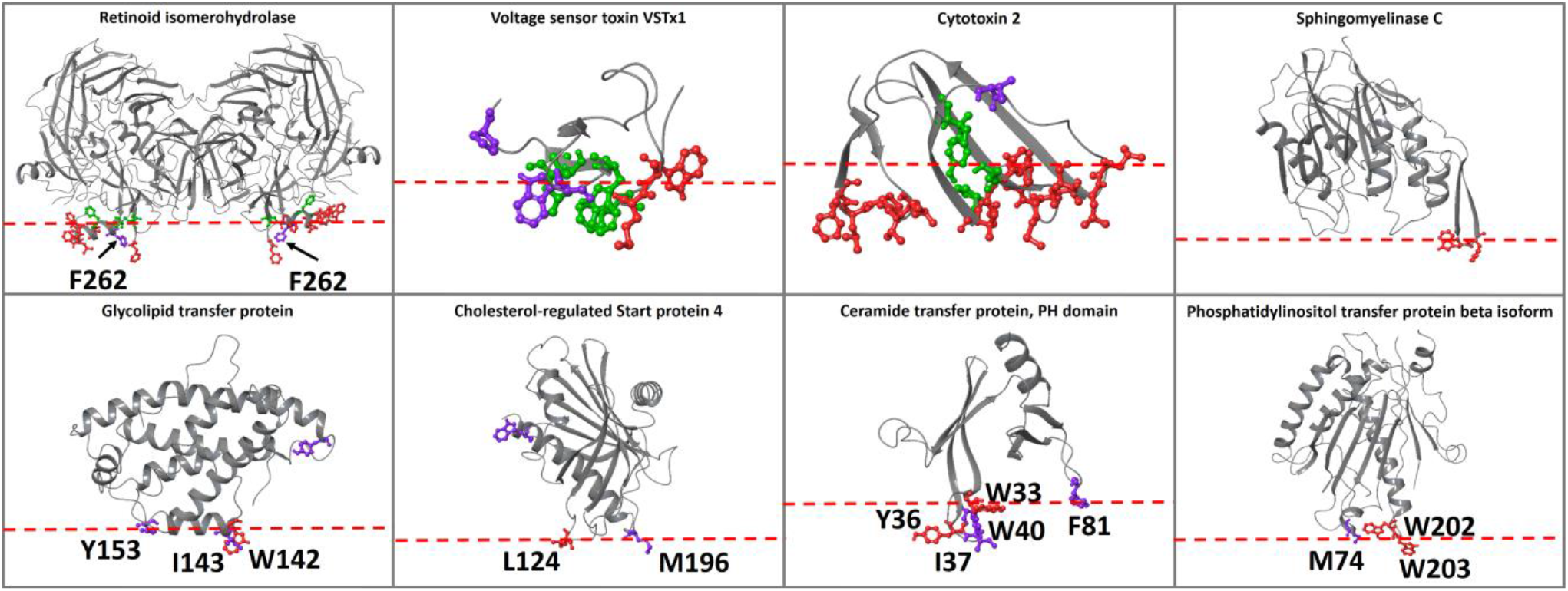
Proteins of the test set. The experimental membrane-penetrating amino acids predicted from the ensemble clas-sifier are depicted in red, the experimental membrane-penetrating amino acids not predicted from the classifier are depicted in green, and the amino acids predicted from the classifier that have not been experimentally-verified are depicted in purple.

Moreover, the ensemble classifier was applied in the test set keeping all amino acid types. To reduce false positive non-hydrophobic amino acids, only those amino acids with center of mass (COM) distance of 14 Å from at least one of the predicted hydrophobic amino acids were kept (Table S9).

Furthermore, the performance of the classifier was compared to two computational tools that predict protein-membrane interfaces from 3D structures, the PPM web-server [46], which additionally predicts the orientation of proteins in membranes, and the MODA web-server [32], in the test set without performing data selection (Table S9). Generally, the predictions of every tool was fairly accurate in predicting the protein-membrane regions with our classifier outperforming them in some cases. Specifically, for the retinoid isomerohydrolase homodimer, PPM falsely predicted the orientation (probably affected by the missing chains) and placed the protein in an orientation in which only one monomer was in contact with the membrane instead of both, while our classifier and MODA correctly predicted the protein-membrane regions in both chains, but with a few false positive predictions for MODA (Figure S3). For the VSTx1 toxin, every tool predicted the protein-membrane region but falsely predicted W25. Moreover, our classifier falsely predicted as membrane-penetrating the amino acids near the N- and C-terminus and MODA falsely predicted the beta sheet on the opposite side of the protein-membrane interface and the C-terminus region (Figure S4). For cytotoxin 2, all tools performed the same with a few false positives for our classifier and MODA in the 41-46 region (Figure S5). Similarly, for sphingomyelinase C all tools recognized the experimentally-verified membrane-penetrating amino acids W284 and F285, with PPM and MODA recognizing amino acids in distant loops that are aligned with the experimentally-verified protein-membrane region suggesting a multiregional interaction with the membrane, which is probably true, according to the proposed membrane binding model (Figure S6). For the glycolipid transfer protein, our classifier and MODA provided similar results correctly identifying the membrane-penetrating α-helix, with MODA falsely predicting the C-terminus and our classifier amino acid Y81 to be membrane-penetrating amino acids. PPM also suggested the insertion of the membrane-penetrating α-helix, but with the addition of the P40-P44 region (Figure S7). Likewise for the cholesterol-regulated Start protein 4, all tools predicted correctly the experimentally-verified amino acid L124 with our classifier and PPM additionally predicting the 196-200 region, MODA falsely predicting the C-terminus, and our classifier falsely predicting amino acid W91 (Figure S8). Finally, for the PH domain of the ceramide transfer protein and the phosphatidylinositol transfer protein beta isoform, the outcome was similar and correct for all tools (Figures S9-S10).

The performance of the ensemble classifier was tested with additional protein use cases with known membrane-penetrating regions (second test set, Figure 3), and the results were compared with PPM and MODA (Table S10). For the cases of cholesterol oxidase, cytochrome P450 3A4, monoglyceride lipase MGLL, L-amino acid deaminase, and intestinal fatty acid binding protein, all tools correctly identified the protein-membrane regions (Figures S11, S12, S14, S19, and S20). For the 9-cis-epoxycarotenoid dioxygenase 1, chloroplastic, all tools predicted the protein-membrane regions, with the exception of our classifier, which predicted the insertion of one of the two parallel amphipathic helices, instead of both (Figure S13). Similarly, for the dihydroorotate dehydrogenase, all tools predicted the protein-membrane regions, with our classifier falsely identifying the amino acid W362 and MODA the region 245-247 (Figure S15). For phosphatase PTEN, our classifier successfully identified the protein-membrane region 263-269 of the C2 domain and the region around the L42 amino acid of phosphatase in the same membrane plane. MODA also identified the same phosphatase region, however it falsely identified the opposite side of the C2 domain as a protein-membrane region. PPM also falsely identified the opposite side of the C2 domain suggesting an orientation, which is opposite to the actual membrane orientation (Figure S16). For (S)-mandelate dehydrogenase, the protein-membrane region was correctly identified by all tools, but our classifier and MODA also identified amino acids 53-56 to be membrane-penetrating (Figure S17); these amino acids are actually at the protein-protein interaction interface in the homotetramer of (S)-mandelate dehydrogenase and are not misclassified if we perform the predictions in the homotetramer biological assembly (Figure S18). For the phosphatidylinositol 4,5-bisphosphate 3-kinase alpha (PI3Kα), all tools predicted amino acids 232-233 as a protein-membrane interface, which is not in agreement with the experimental results, but is in fact the region where PI3Kα binds to K-RAS (Ras binding domain). Additionally, our classifier and MODA successfully identified the p110α 863-872 and the iSH2 512-525 regions, but falsely identified the 498-508 region, which links the C2 domain with the helical domain (Figure S21). The membrane orientation resulting from PPM is different from the one proposed through mutagenesis experiments [14]. Finally, for the phosphatidylcholine transfer protein, all tools provided the same results identifying the experimentally proven membrane-interacting region 184-193 and an adjacent loop, while MODA additionally predicted loop 147-148 to be membrane binding, which is in the same plane with the other two membrane binding regions (Figure S22).

**Figure 3.**
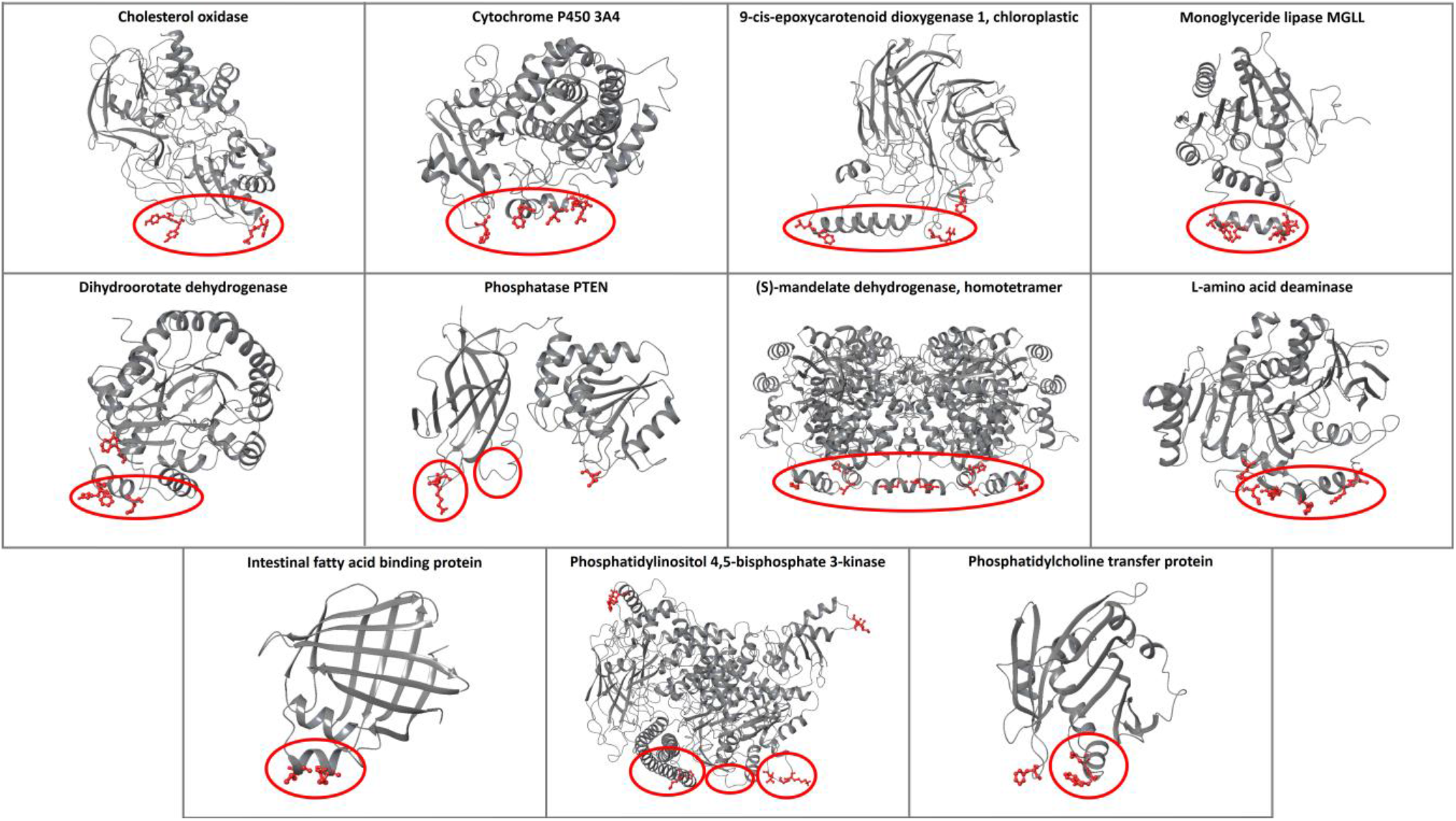
The predictions of the ensemble classifier for proteins with known protein-membrane interface regions. The membrane-penetrating amino acids predicted from the classifier are depicted in red, and the experimental membrane-penetrating regions are denoted with red circles.

Furthermore, the ensemble classifier was applied in the full structure of prothrombin (PDB: 5EDM [47]) and the results of identified membrane-protein interfaces were compared with PPM and MODA (Figure 4). All tools predicted the GLA domain 3-5 region as a membrane contacting region, which is natural for the classifier because the GLA domain of prothrombin was in the training set. Additionally, the classifier predicted the amino acids Y93, W398, and V458 and MODA predicted the amino acids Y93, Y377, R379, and R484, suggesting an orientation parallel to the actual membrane orientation, opposed to the perpendicular suggested by PPM. Y93 is a key prothrombin amino acid, which is essential for stabilizing the closed form and protects the active site pocket of the protease domain [48]. In the prothrombin closed form (PDB: 6BJR [48]), Y93 inserts its aromatic side chain into the binding pocket of the protease domain engaging W547 (W533 of 5EDM) and forms pi-pi interactions (Figure S23). The results provided by our classifier and MODA suggest that Y93 penetrates into the membrane, further indicating that when prothrombin engages the membrane the open form is favored with Y93 anchoring the membrane and opening the active site.

**Figure 4.**
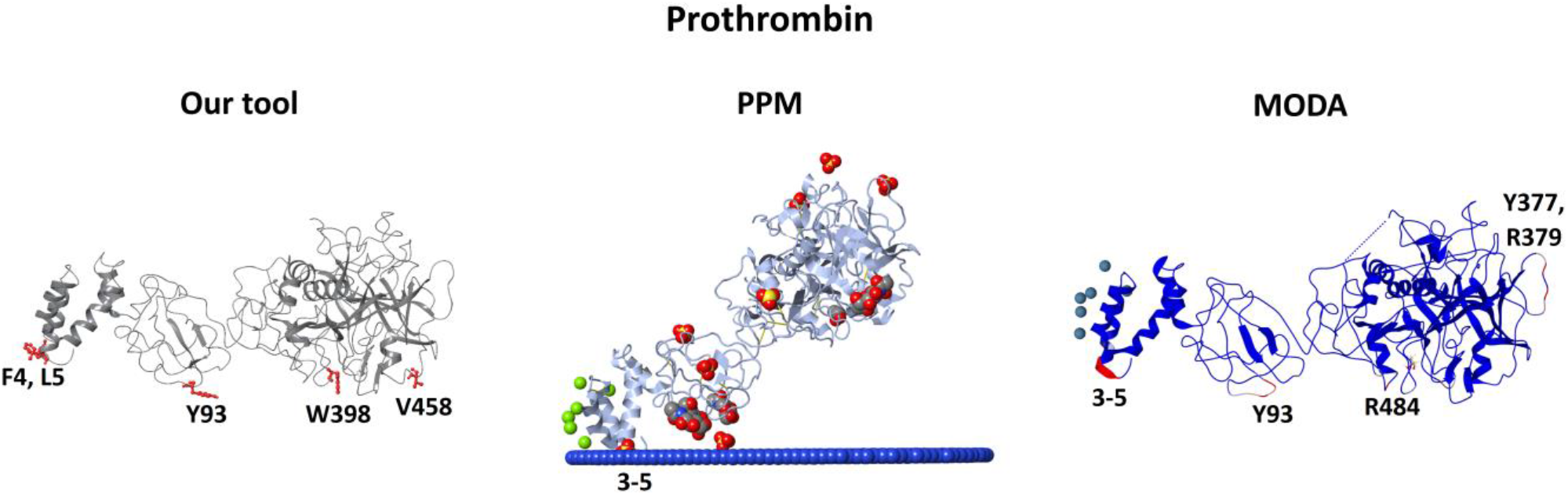
Comparison of the predictions provided from our classifier, PPM, and MODA for the open form of the pro-thrombin protein. Our classifier and MODA propose a parallel to the membrane orientation, suggesting the in-sertion of Y93 to the membrane, which in turn opens the active site of the protease domain (Figure S23).

Finally, the ensemble classifier was also tested for the prediction of the protein-membrane interfaces of nine transmembrane enzymes described in Ref [49], which include a soluble domain performing extracellular catalysis. In agreement with experimental results, our classifier predicted numerous amino acids that lie in the hydrophobic lipid bilayer core, along with membrane-interacting extracellular amino acids (Figure S24).

## Discussion and Conclusions

Drug design of protein-membrane interfaces for peripheral membrane proteins has been so far neglected due to the complexity of the interface and the lack of a suitable workflows and simulation technology capable of implementing this drug design strategy. Furthermore, protein-membrane interaction regions of peripheral membrane proteins are commonly unknown, and only a few rational methodologies exist that predict these regions from the 3D protein structure. To assist in protein-membrane interface recognition, a novel ensemble machine learning classifier is described trained in experimental data retrieved from extensive literature search.

The ensemble classifier results are accurate in predicting correctly the membrane-penetrating amino acids in the test set, providing with a macro-averaged F_1_ score = 0.92 and an MCC = 0.84. Additionally, in a different independent dataset with experimentally known protein-membrane regions, our classifier correctly identified membrane-penetrating amino acids in these regions with a few false positive predictions. In addition, comparative results demonstrated that our classifier performed similarly, and in some cases better, than the only two available web-servers that predict protein-membrane interaction sites from the 3D protein structure, PPM and MODA. Moreover, our classifier successfully predicted the membrane-penetrating amino acids and the amino acids that lie in the hydrophobic core of the lipid bilayer in transmembrane proteins containing a soluble catalytic domain.

The features used in this study have a significant impact on the performance of the ensemble classifier. The fact that except from the hydrophobicity and the solvent exposure, other features—such us evolutionary conservation, secondary structure, flexibility, dihedral angles, and TAE descriptors—are important in our model decision-making process, offers novel physicochemical insights in the way that peripheral membrane proteins attach to the membrane.

Commonly, during development of computational tools several obstacles may emerge. In this case the first obstacle was the low number of the peripheral membrane proteins with experimentally known membrane-penetrating amino acids described in literature. The second and more crucial constraint was the small number of amino acids that were tested experimentally resulting in membrane-penetrating amino acids being marked as non-penetrating, which in turn resulted in misinforming the classifiers during the training process and rendering the selection of the best ensemble classifier strenuous. Moreover, based on the fact that membrane-penetrating amino acids are labeled as non-penetrating, the performance metrics (e.g. F scores) do not reflect the actual accuracy of the ensemble classifier, which is higher, and therefore, a direct numerical comparison of our classifier, PPM, and MODA results is not possible.

Manual inspection of false positive results revealed that several amino acids were located near the N- or C-terminus, or near missing loops, probably because the area is more solvent exposed. Intriguingly, other amino acids falsely predicted as membrane-penetrating are found to be implicated in protein-protein interactions. For example, in the case of (S)-mandelate dehydrogenase our model correctly classifies membrane-penetrating amino acids in the homotetramer form, while in the homodimer form we find that amino acids that are implicated in protein-protein interactions are also classified as membrane-penetrating. Hence, it is advisable to use the complex protein structure if it is available. The assumption that protein-membrane interactions are similar to protein-protein interactions was also deduced by [32], who adapted their protein-protein interaction interface prediction PIER algorithm [33] in MODA.

Also, it should be noted that the predictions depend on structural information, therefore, in the case where in the 3D protein structure the membrane-penetrating amino acids are in the bulk of the protein and a conformational change is necessary to face them towards the membrane, or the protein is intrinsically disordered, the ensemble classifier would not be able to predict them. Furthermore, it should be noted that neither our model, nor PPM or MODA are suitable tools for classifying if a protein is a peripheral membrane protein or not. A sequence-/evolutionary-based deep learning classifier would be more appropriate for this purpose [50]. Also, with the recent advancements in protein structure predictions, i.e., AlphaFold2 [51] and RoseTTAFold [52], the structure of unresolved proteins can be predicted with high accuracy, but in many cases, these models fail to fold the N- or C-terminus, or various protein segments. It is recommended to remove these regions, i.e., amino acids with confidence score (pLDDT) less than 70, before applying our model, since these unfolded regions are going to affect the prediction accuracy.

In the future, we plan to devise methods that orient peripheral membrane proteins in the membrane. Such an approach could include (1) placing the protein in a model membrane in all possible orientations based on the ensemble classifier predictions, (2) measuring the protein-membrane interface energy with an energy function specific for this purpose, and (3) retain the protein-membrane orientation with the lowest energy. When an appropriate method to define the membrane-protein orientation is devised, designing a web-database with our model predictions, similar to PPM and OPM [46] could also be envisaged.

Moreover, modifying the labeling system by dissecting the structures by a well-defined plane into membrane penetrating and not penetrating parts, or by labeling any amino acid from a specific distance from the experimentally known membrane penetrating amino acids as a true label warrants further investigation. Such a consideration would improve the class imbalance problem, although a defined plane and distance may be subjective to generate.

Finally, it is noteworthy to mention that the membrane-penetrating amino acids are in many cases significant for the allosteric control of binding sites similar to prothrombin case. It is suggested that in the protein-membrane interfaces a binding pocket exists [18, 20–25]. We strongly believe that these binding pockets could act allosterically being connected with the active site and could be able to modulate the open/closed form equilibrium [17, 53] or even block protein function by disrupting the protein-membrane interactions.

## Key points

- A dataset of peripheral membrane proteins with experimentally known membrane-penetrating amino acids was assembled.
- An ensemble machine learning classifier was trained utilizing thermodynamic, topographic, and property-based features.
- The ensemble machine learning classifier displayed a macro-averaged F_1_ score = 0.92 and an MCC = 0.84 in identifying membrane-penetrating amino acids.
- The python code is publically available for usage at https://github.com/zoecournia/DREAMM.

## Supplementary data

Supplementary data are available online at https://academic.oup.com/bib.

## Funding

This work was supported by the State Scholarships Foundation (ΙΚΥ) [MIS-5000432 to A.C.]; the Hellenic Foundation for Research and Innovation (H.F.R.I.) [1780 to Z.C.]; and the high-performance computing resources provided by the National Infrastructures for Research and Technology S.A. (GRNET S.A.) [pr008033_gpu-Mem-Surf, pr008033_thin-Mem-Surf, and pr009022_gpu-Mem-Surf].

## Conflict of Interest

The authors declare no conflict of interest.

## Notes

https://github.com/zoecournia/DREAMM

### Competing Interest Statement

The authors have declared no competing interest.

