## Supplementary information for "Predicting protein-membrane interfaces of peripheral membrane proteins using ensemble machine learning"

### Feature extraction

Lipid interfaces of proteins possess specific chemical and topological properties, *i.e.*, amphipathic alpha-helices flanked by a flexible hinge or loop regions, being solvent-exposed or containing cationic patches around aromatic and aliphatic regions that anchor to the negatively-charged bilayers [1, 2], which are regularly found in the inner leaflet of the plasma membrane [3]. Therefore, two driving forces for protein-membrane association need to be considered: a) long range electrostatic interactions that drive protein-membrane proximity, b) hydrophobic interactions that facilitate protein anchoring to the hydrophobic fatty acid tails of the lipid bilayer [2, 4].

Firstly, the ProtDCal tool was facilitated, which calculates 2788 thermodynamic, topographic, and property-based features [5], and then, based on the protein-membrane interactions, physicochemical and biochemical features of the aforementioned membrane-associated properties were generated for each protein amino acid in the training set by employing computational methods in Python as described below. Protein-membrane regions mainly consist of amphipathic alpha-helices or hydrophobic loops, therefore, the Define Secondary Structure of Proteins (DSSP) program was utilized to define the secondary structure [6], parsing the PDB files with the Python package Biopython [7, 8]. Additionally, DSSP measures geometrical properties, for example, backbone torsion angles, which were also kept as features. One-hot encoding, a technique that transforms each unique value in a categorical feature into a new binary feature, was applied on the amino acid and the secondary structure features to transform them from categorical to numerical features. Alongside the secondary structure, the solvent exposure is a significant property of the membrane-penetrating amino acids. The FreeSASA tool was utilized for the calculation of the SASA [9], and the MSMS tool for calculating the amino acid and Cα depths [10]. One advantage of FreeSASA lies in the separation of amino acid SASA into polar and non-polar SASA. Non-polar SASA combines solvent exposure with hydrophobicity, which is necessary for forming hydrophobic interactions with the inner leaflet of the plasma membrane. Therefore, the Wimley-White whole-residue interface and octanol hydrophobicity scales were additionally utilized as features [11, 12]. To consider electrostatic interactions, PDB2PQR was utilized to calculate amino acid charges and protonation states using default parameters [13, 14], and MDAnalysis for reading the resulting PQR file [15].

Moreover, additional properties (features), which could potentially be connected to the protein-membrane association, were sought. The sequence profiling tool HHblits was applied in order to calculate the conservation score [16], through HTMD [17], searching the Uniclust30 database [18]. To consider amino acid flexibility, the ProDy package was used to calculate squared fluctuations utilizing two different Elastic Network Models, the Gaussian Network Model and the Anisotropic Network Model [19, 20]. Finally, the feature space was enriched with the amino acid radius of gyration, which was calculated with MDAnalysis [15], the number of each amino acid type of neighboring amino acids, and their total number in a Cα – Cα distance of 7 Å.

To consider the surrounding amino acid properties of each amino acid, the mean values of the aforementioned features were calculated, for each amino acid and the amino acids at a 7 Å distance from the protein α carbon atoms (Cα – Cα). In this way, the 3D space is taken into consideration ensuring that the information of the surrounding amino acids is included in the feature space, for example, the mean hydrophobicity and charge from the neighboring amino acids, and the number of nearby lysine, arginine, and histidine amino acids, leading to 2880 features in total. Feature and data selection followed to discard redundant information as it is discussed in the SI, and the feature importance was calculated with the most important features expressing hydrophobicity and solvent exposure (Figure S1, Table S3).

**Feature and data selection**

Training machine learning algorithms in datasets with redundant samples and features is computationally inefficient. Discarding redundant information is essential to reducing the data size and hence the computational effort as in this case the training set consists of 12,805 samples (amino acids) and 2,880 features. To reduce the sample size, and especially the sample size of the majority class, the solvent inaccessible amino acids of the proteins were removed because only interfacial amino acids penetrate the membrane, using a cutoff of 2.5 Å for the amino acid depth feature produced by MSMS [10], leading to 8,720 samples. Moreover, only amino acids that penetrate the hydrocarbon core of the membrane were retained, according to the octanol-interface scale [21], which is derived from the experimentally determined Wimley-White whole-residue interface and octanol hydrophobicity scales [11, 12], leaving out the A, S, N, G, E, D, K, R, and H amino acids, further reducing the sample size of the majority class. The Wimley-White whole-residue hydrophobicity scales are experimentally determined transfer free energies ΔG (kcal/mol) from water to POPC interface ($\Delta G_{IF}$) and to n-octanol ($\Delta G_{oct}$). From the experimentally-verified membrane-penetrating amino acids, we included in the membrane-penetrating class the amino acids that are hydrocarbon-preferring ($|\Delta G_{oct} - \Delta G_{IF}| \leq 0.25$ kcal/mol) [21]. The C and Q amino acids were also removed, as only a few cases were present in the membrane-penetrating category. In the end, the training set was reduced to 3010 samples, reducing the required computational time to process the dataset. Moreover, the class imbalance problem was attenuated because the sample size of the majority class was reduced significantly, with the membrane-penetrating amino acids consisting the ~5.5% of the total samples.

Subsequently, features with zero standard deviation (all feature values are the same) were discarded in order to remove redundant features, further reducing the dataset to 2252 features. The Pearson pairwise correlation was measured leaving out features with more than 95% correlation; this led to 883 features. Utilizing two different tree-based machine learning algorithms, the extremely randomized trees method [22], using the scikit-learn Python package [23] and the leaf-wise gradient boosting decision tree algorithm LightGBM [24], features that where less important than the 1% of the total feature importance in both algorithms were removed leading to a total of 727 features. The hyper-parameter space for these classifiers was fine-tuned by combining the randomized and grid search 5-fold cross validation approaches as explained in the "Ensemble machine learning methodology" section in the main text. As anticipated, the most important features were expressing hydrophobicity and solvent exposure (Figure S1, Table S3). Finally, the features were again inspected resulting in 167 redundant features that do not affect the behavior of the newly developed ensemble classifier and leading to 560 features, accelerating the feature extraction process.

**Class imbalance problem**

Reducing the data sample size improved the class imbalance problem but the increase of the membrane-penetrating class to almost five-fold still produced imbalanced classes. When the size of one class outnumbers the size of the other, machine learning algorithms will under-predict the infrequent class. To balance the two classes three techniques were utilized. The first one is to use weights on the samples, emphasizing the minority samples. The second is to over-sample the minority class using algorithms that generate synthetic samples based on the feature values of the minority class samples until both classes consist of equal number of samples. Finally, the third technique is to under-sample the majority class with sample selection methods until again both classes have equal number of samples. Notably, for a number of machine learning algorithms using weights is similar to over-sampling the minority class with duplicate samples. From the initial training set of 3,010 samples, two training sets of 5,686 samples using two different over-sampled techniques, two training sets of 334 samples using two different under-sampled techniques, and one more training set of 4,287 samples using a combination of over- and under-sampling methods were produced utilizing the imbalanced-learn Python toolbox [25]. Using this procedure, six different training sets were produced: the initial training set using weights, two training sets using over-sampled techniques, two training sets using under-sampled techniques, and one training set using a combination of over- and under-sampling methods.

The first over-sampled training set was generated utilizing the Synthetic Minority Over-sampling Technique (SMOTE) technique, which synthesizes artificial new minority instances between existing real minority instances [26]. The second over-sampled training set utilizing the Adaptive Synthetic (ADASYN) sampling algorithm is similar to SMOTE but attempts to infer which points in the minority class would be the most difficult for a model to learn and attempts to place a higher ratio of synthetic data close to these points [27]. For the under-sampled training sets, the first one was generated using the Condensed Nearest Neighbor (CNN) method, which iteratively uses the 1 nearest neighbor rule to decide if a sample should be removed or not [28] and the second one was based on the Instance Hardness Threshold (IHT), which is a technique where a machine learning algorithm is trained on the training set and removes the samples with the lowest probabilities [29]. For the IHT method the scikit-learn gradient boosting classifier [30] was utilized to estimate the instance hardness of the samples. Because over-sampling using SMOTE may lead to generation of noisy samples, a sixth training set was built, where SMOTE was followed by the Edited Nearest Neighbors undersampling method (SMOTEENN), which applies a nearest-neighbors algorithm to clean the training set [31]. By removing samples which do not agree enough with their neighborhood (the majority of the 5 closest neighbors belong to another class) the outliers that were generated from SMOTE were removed. These six training sets can be visualized in Figure S2 reduced to two dimensions.

**Performance metrics**

To measure the performance of machine learning classifiers, the number of correct and incorrect predictions is considered. We can summarize the predictions with count values for each class (here: membrane penetrating amino acids and non-membrane penetrating) into a matrix. This matrix is called the confusion matrix, and for binary classifications problems is derived as:

***Confusion Matrix***

| | | True class | |
|---|---|---|---|
| | | **Positive** | **Negative** |
| **Predicted class** | **Positive** | **TP** | **FP** |
| | **Negative** | **FN** | **TN** |

where TP stands for true positives, FP for false positives, FN for false negatives, and TN for true negatives. The performance metrics for a machine learning binary classifier are computed from the confusion matrix. The most common performance metric is accuracy (eq. 1), which is the fraction of correct predictions over the total predictions:

$$Accuracy = \frac{TP + TN}{Total\ predictions} \quad (1)$$

The accuracy score metric may be misleading in imbalanced datasets. For example, consider that the number of samples of the minority class (e.g. positive class) is 100 and the number of samples of the majority class (negative class) is 9,900. Then, if the algorithm classifies all samples to be in the negative class, then the confusion matrix becomes:

***Confusion Matrix***

| | | True class | |
|---|---|---|---|
| | | **Positive** | **Negative** |
| **Predicted class** | **Positive** | **TP=0** | **FP=0** |
| | **Negative** | **FN=100** | **TN=9,900** |

and the accuracy score is derived as,

$$Accuracy = \frac{0 + 9{,}900}{10{,}000} = 0.99$$

which does not represent the actual performance of the model as the 100 samples of the minority class were wrongly labeled in their entirety. To overcome this limitation of the accuracy score, more sophisticated metrics have been introduced. For example, the $F_1\, score$ is derived by taking the harmonic mean of the precision metric and the recall metric (see main text). The $F_1\, score$ measures the performance in each class separately, and the macro $F_1\, score$ (eq. 2) is the average performance of all classes with values ranging between 0 and 1:

$$Macro\ F_1\ score = < F_1\ score > \quad (2)$$

where angle brackets denote average over each class' $F_1\, score$.

Another metric is the Matthews correlation coefficient (MCC) (eq. 3). The MCC metric receives values between -1 and 1 and is formulated as,

$$MCC = \frac{TP * TN - FP * FN}{\sqrt{(TP + FP)(TP + FN)(TN + FP)(TN + FN)}} \quad (3)$$

which is an alternative metric for measuring the performance of binary classifications. In the previous example, the $macro\ F_1\ score = \frac{0+0.95}{2} = 0.48$ and $MCC = 0$, correctly measuring the performance.

Herein, except from deriving the harmonic mean of precision and recall ($F_1\, score$), the geometric mean of sensitivity (eq. 4) and specificity (eq. 5) is also measured is also measured. The sensitivity and specificity are formulated as:

$$sensitivity = \frac{TP}{TP + FN} \quad (4)$$

$$specificity = \frac{TN}{TN + FP} \quad (5)$$

and the geometric mean (eq. 6) of the specificity and sensitivity is formulated as:

$$geometric\ mean\ score = \sqrt{\text{sensitivity} * \text{specificity}} \quad (6)$$

And similarly to the $F_1\, score$, the macro average is taken.

Another metric to measure the performance of the machine learning models is area under the curve (AUC) of the precision-recall curve (the precision-recall curve is the plot of precision vs recall). For example, if the samples of the minority class (e.g. positive class) are 100 and the samples of the majority class (negative class) are 9900, then the classifier predictions are summarized in the following confusion matrix:

***Confusion Matrix***

| | | **True class** | |
|---|---|---|---|
| | | **Positive** | **Negative** |
| **Predicted class** | **Positive** | **TP=90** | **FP=900** |
| | **Negative** | **FN=10** | **TN=9000** |

the values of precision and recall are 0.09 and 0.9, respectively, and the precision-recall AUC is 0.5, correctly capturing the classifier behavior.

**Hyper-parameter optimization**

Each machine learning classifier has parameters that control the behavior of the training algorithm and impact the performance. These parameters are called hyper-parameters and affect the classifier capability of identifying patterns and correlations. For example, hyper-parameters may be the number of neighbors in k-nearest neighbors, the kernel type in support vector machines, the maximum depth of the tree in a decision tree, the number of trees in random forests, etc. A description for each hyper-parameter for each classifier employed herein may be accessed in https://scikit-learn.org/stable/modules/classes.html, https://xgboost.readthedocs.io/en/latest/parameter.html, and https://lightgbm.readthedocs.io/en/latest/Parameters.html. It is infeasible to know a-priori the optimal hyper-parameters for each classifier. Therefore, a randomized search cross-validation technique was performed in a wide range of values for each hyper-parameter set (Table S6), training hundreds of thousands of models. The best hyper-parameter sets based on the randomized search cross-validation were selected and then the grid search cross-validation technique was used in the vicinity of this best hyper-parameter space to search again for the optimal hyper-parameters (Table S6).

Each hyper-parameter combination was assessed by applying the 5 fold cross-validation using the $F_1\ score$; the training set was split in 5 folds, training with the 4 folds and validating with the remaining fold. This was repeated 5 times, each time with a different fold as a validation set, and in the end, the average of the 5 $F_1\ scores$ was calculated. This procedure was repeated for each hyper-parameter combination. The hyper-parameter combination with the best average $F_1\ score$ was kept. For assessing the meta-classifiers, the $F_2\ score$ (eq. 7) was chosen to emphasize on recall.

$$F_2\ score = 5 * \frac{precision * recall}{(4 * precision) + recall} \quad (7)$$

**Percentage Identity**

To ensure that the test sets are unbiased, the fold and family of each protein was retrieved from "Structural Classification of Proteins" (SCOP) web-database [32, 33], and the percentage identity between all sequences of the dataset (training and test sets) was calculated. Firstly, a local sequence alignment was performed for every pair of sequences utilizing the Biopython Python package [34]. The gap opening and gap extension penalty values were chosen to be -10 and -1, respectively. The local alignment was preferred over the global alignment because it better captures conserved motifs and domains in divergent sequences. Then, the percentage identity (eq. 8) was calculated for every aligned pair with the following formula:

$$Percentage\ identity = \frac{Identical\ positions}{\min(TG_A, TG_B)} \quad (8)$$

Where $TG_A$ and $TG_B$ are the sum of the number of amino acids and internal gap positions in sequences A and B in the alignment as in Ref. [35].

High percentage identity values (more than 40%) were observed for some proteins in the training set, but not in the test sets (Table S5), ensuring that the test sets predictions are unbiased. To better observe biased data, the CD-HIT Suite [36] web-server was employed in the whole dataset to capture clusters of similar proteins, choosing the minimum available sequence identity cutoff (40%). The results demonstrated five clusters of similar sequences, again, with no protein sequences from the two test sets (Table S4).

**Table S1.** The peripheral membrane proteins of the training set, their PDB code, their experimentally known hydrophobic membrane-penetrating amino acids after data selection, and the experimental method used to determine them. Amino acid numbering is the same as in the PDB structures.

| Protein | PDB | Membrane-penetrating amino acids | Methods & References |
|---|---|---|---|
| Cytosolic phospholipase A2, group IVA | 1cjy, chain A | F35, M38, L39, Y96, V97, M98, W464 | Bn, SL [37-40] |
| Synaptotagmin-1, C2A domain | 2k45 | M34, F95 | Bn, SL [41, 42] |
| Protein kinase Cα, C2 domain | 4dnl | P188, T250, T251 | XRR, NMR [43, 44] |
| Synaptotagmin-1, C2B domain | 4v11 | V305, I368 | SL [45] |
| Protein kinase C epsilon, C2 domain | 1gmi | W23, I89, Y91 | Bn [46] |
| Neutrophil cytosol factor 4 (p40phox) | 1h6h | F35, Y94, V95 | Bn [47] |
| Neutrophil cytosol factor 1 (p47phox) | 1kq6 | I65, W80 | Bn [47, 48] |
| Early endosome antigen 1 FYVE domain | 1joc | V1367, T1368, V1369 | NMR [49, 50] |
| Vps27 FYVE domain | 1vfy | L185, L186 | Bn [51] |
| C1 domain of protein kinase C delta | 3uej, chain A | M239, P241, T242, F243, L250, W252, L254, V255 | Bn, NMR [52] |
| Epsin ENTH domain | 1h0a | L6, M10, I13, V14 | Bn [53] |
| C2 domain of coagulation factor V | 1czs | W26, W27 | Bn [54] |
| C2 domains of coagulation factor Va | 1sdd | Y1943, L1944, W2050, W2051 | Bn [55] |
| Coagulation factor VIII | 2r7e | W2093, M2199, F2200, L2251, L2252 | Bn [56, 57] |
| Annexin V | 1anx, chain A | T74, W187 | Bn [58] |
| Equinatoxin II | 1iaz, chain A | W112, Y113 | Bn, NMR, Fn [59-61] |
| Prostaglandin H2 synthase 1 | 2ayl | I74, W75, W77, L78, F88, F91, L92, W98, L99, F102 | Bn, Fn [62] |
| Pancreatic phospholipase A2, group IB | 1hn4, chain A | W3, L19, M20 | FLq [63] |
| Bee venom phospholipase A2 | 1poc | I1, I2, F24, I78, F82 | FLq [64] |
| Phospholipase A2, group IIA | 5g3n, chain A | L2, V3, L19, F23, V30, F63 | FLq [64] |
| Acidic phospholipase A2 1 | 1poa | Y3, W19, W61, Y63, F64, Y110 | Bn, HD [65-67] |
| Snake phospholipase A2, group II | 1vap, chain A | W20, W30, W109 | FL [68] |
| Snake phospholipase A2, group II, B | 4hg9, chain A | Y120, P121, I124, L125 | Bn [69] |
| Phosphatidylinositol-specific phospholipase C | 2ptd | I43, W47, W242 | Bn [70] |
| α-toxin (bacterial phospholipase C) | 1gyg, chain A | Y331, F334 | Fn [71] |
| Arachidonate 15-lipoxygenase | 2p0m, chain A | Y15, F70, L71, W181, L195 | Bn [72] |
| 8R-Lipoxygenase | 2fnq, chain A | W413, F414, Y448, W449 | FL, Fn [73, 74] |
| Signal peptidase I | 3iiq, chain A | W300, W310 | Fn, FLq [75, 76] |
| Perfringolysin | 1pfo | W466, T490, L491 | Bn [77, 78] |

| | | | |
|---|---|---|---|
| Hepatocyte growth factor-regulated tyrosine kinase substrate (Hrs) | 1dvp | F173 | Bn [51] |
| Prothrombin, GLA domain | 1nl1 | F5, L6, V9 | FLq [79] |
| Sorting nexin-3 | 5f0p | From chain C F103, F110 | NMR [80] |
| Beta-2-glycoprotein 1 | 1c1z | L313, F315, W316 | Bn [81] |
| Alpha-tocopherol transfer protein | 1oiz, chain A | F165, F169, I202, V206, M209 | Bn [82] |
| Annexin 24 | 1dk5, chain A | W35, W107, Y192 | FL [83] |
| Protein kinase C gamma type C1B domain | 1tbn | Y123, L125 | Fn [84] |
| Protein kinase C gamma type C1A domain | 2e73 | W57, I59 | Fn [84] |
| RAF-1 proto-oncogene serine/threonine-protein kinase | 1faq | L147, L149, F158 | NMR [85] |
| Eosinophil cationic protein | 4x08, chain A | W35 | NMR [86] |
| Antimicrobial peptide kalata B1 | 2mh1 | W22, P23, V24, L30, P31, V32 | NMR [87] |
| Kappa-theraphotoxin-Scg1a | 1la4 | Y4, L5, F6, W30 | Fn, FLq [88, 89] |
| Dual adapter for phosphotyrosine and 3-phosphotyrosine and 3-phosphoinositide | 1fao | L177, V178 | Bn [90] |
| Pleckstrin homology domain-containing family A member 1 | 1eaz | V204, M205 | Bn [90] |
| Sticholysin II | 1gwy, chain A | W110, Y111, W114, Y136 | Fn [91, 92] |
| Sticholysin I | 2ks4 | F51, F107, Y109, W111, Y112, W115, M135, Y136, Y137 | NMR [93] |
| Matrix protein VP40 | 1es6 | L295, V298 | Bn, FLq [94, 95] |

Bn – based on membrane binding affinities of mutants; SL – spin-labeling data based on depth parameters Φ; NMR – chemical shift perturbation; FL – fluorescence; FLq – fluorescence quenching; Fn – functional studies of mutants; XRR – X-ray reflectivity; HD – hydrogen-deuterium exchange mass spectrometry.

**Table S2.** The peripheral membrane proteins of the test set 1, their PDB code, their experimentally known hydrophobic membrane-penetrating amino acids after data selection, and the experimental method used to determine them. Amino acid numbering is the same as in the PDB structures.

| Protein | PDB | Membrane-penetrating amino acids | Methods & References |
|---|---|---|---|
| Retinoid isomerohydrolase | 3fsn | F196, F200, I202, F264, L265, W268, L270, W271 | X-ray [96] |
| Voltage sensor toxin VSTx1 | 1s6x | F5, M6, W7, W27, V29, L30 | FLq, FL, NMR [97-99] |
| Cytotoxin 2 | 1ffj | L6, V7, P8, L9, F10, Y22, M24, F25, M26, V27, P30, V32, P33, V34, I39, L47, L48, V49 | NMR [100] |
| Sphingomyelinase C | 2ddr, chain A | W284, F285 | Bn [101] |
| Glycolipid transfer protein | 3rzn | W142 | Fn, NMR [102, 103] |
| Cholesterol-regulated Start protein 4 | 1jss, chain A | L124 | Bn [104] |
| Ceramide transfer protein, PH domain | 2rsg | W33, Y36 | Fn, NMR [105] |
| Phosphatidylinositol transfer protein beta isoform | 2a1l | W202, W203 | Bn [106] |

Bn – based on membrane binding affinities of mutants; NMR – chemical shift perturbation; FL – fluorescence; FLq – fluorescence quenching; Fn – functional studies of mutants; X-ray – X-ray crystallography.

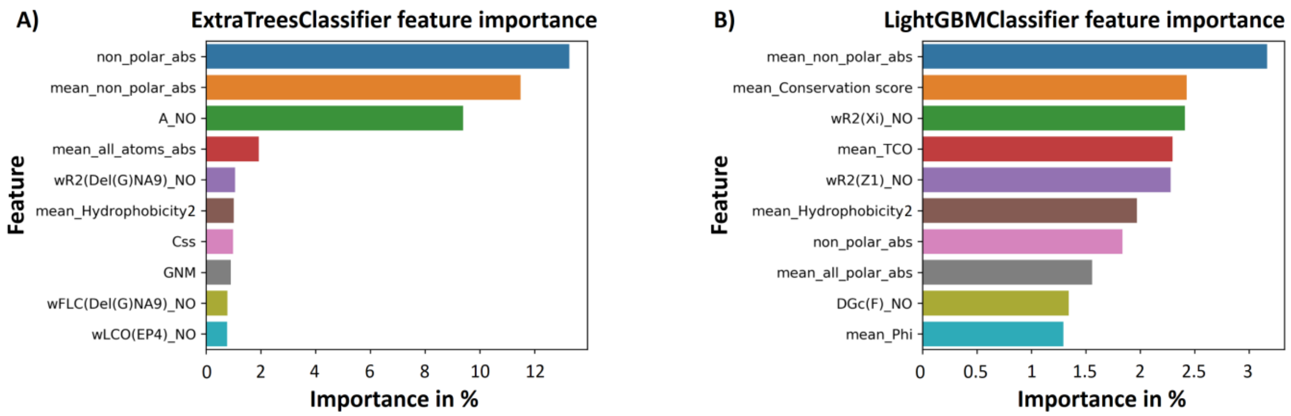


**Figure S1.** The 10 most important features for A) extremely randomized trees classifier and B) LightGBM classifier.

**Table S3.** Description of the most important features from the two tree-based approaches. The most important features express hydrophobicity and solvent exposure. The mean values are for each amino acid and the surrounding amino acids in a Cα – Cα distance of 7 Å.

| Amino acid Feature Name | Description |
|---|---|
| non_polar_abs | Non polar SASA |
| mean_non_polar_abs | Mean non polar SASA |
| A_NO | SASA |
| mean_all_atoms_abs | Mean SASA |
| wR2(Del(G)NA9)_NO | Weighted squared mean radius of the gradient of electronic kinetic energy density perpendicular to the atomic surface [107] |
| mean_Hydrophobicity2 | Mean Wimley-White hydrophobicity scale from water to n-octanol |
| Css | Coil loop secondary structure |
| GNM | Squared fluctuations calculated with GNM |
| wFLC(Del(G)NA9)_NO | Weighted fraction of local contacts of the gradient of electronic kinetic energy density perpendicular to the atomic surface [107] |
| wLCO(EP4)_NO | Weighted amino acid local contact order of an electrostatic potential descriptor [107] |
| mean_Conservation_score | Mean conservation score |
| wR2(Xi)_NO | Weighted squared mean radius of the torsional angle ξ [108] |
| mean_TCO | Mean cosine of angle between C=O of amino acid i and C=O of amino acid i-1 |
| wR2(Z1)_NO | Weighted squared mean radius of a combined measure of hydrophobicity related properties [109] |
| mean_all_polar_abs | Mean polar SASA |
| DGc(F)_NO | Index of the contribution to the free energy from the conformational entropy in a folded state |
| mean_Phi | Mean phi backbone torsion angle |

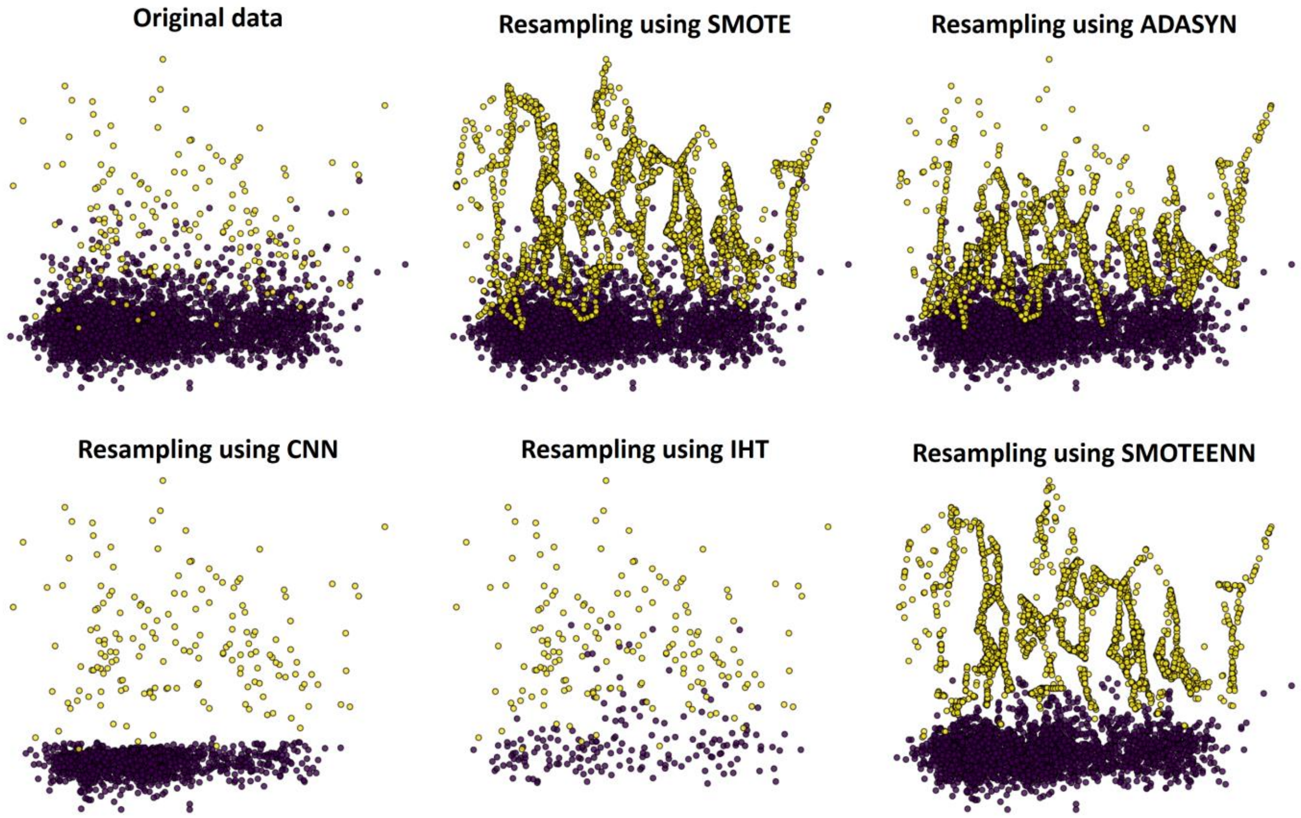


**Figure S2.** A visual representation of our six training sets reduced to two dimensions. The x-axis is the first principal component from principal component analysis (PCA) and the y-axis the component from linear discriminant analysis (LDA).

**Table S4.** The results of CD-HIT Suite web-server [36] using as input the whole dataset and a 40% sequence identity cutoff.

| Cluster | Protein | PDB | Sequence identity |
|---|---|---|---|
| 0 | Pancreatic phospholipase A2, group IB | 1hn4, chain A | Representative |
| | Acidic phospholipase A2 1 | 1poa | 55.93% |
| | Snake phospholipase A2, group II | 1vap, chain A | 41.46% |
| | Phospholipase A2, group IIA | 5g3n, chain A | 40.94% |
| | Snake phospholipase A2, group II, B | 4hg9, chain A | 40.16% |
| 1 | Sticholysin I | 2ks4 | Representative |
| | Sticholysin II | 1gwy, chain A | 93.14% |
| | Equinatoxin II | 1iaz, chain A | 65.71% |
| 2 | Hepatocyte growth factor-regulated tyrosine kinase substrate (Hrs) | 1dvp | Representative |
| | Vps27 FYVE domain | 1vfy | 43.28% |
| 3 | Protein kinase C gamma type C1B domain | 1tbn | Representative |
| | C1 domain of protein kinase C delta | 3uej, chain A | 49.23% |
| 4 | C2 domains of coagulation factor Va | 1sdd | Representative |
| | C2 domain of coagulation factor V | 1czs | 89.38% |

**Table S5.** The peripheral membrane proteins of the dataset, their PDB code, their fold and family, and the sequence identity percentage with the PDB resulting the biggest percentage.

| Protein | PDB | Fold | Family | Percentage identity |
|---|---|---|---|---|
| **Training set** | | | | |
| Cytosolic phospholipase A2, group IVA | 1cjy, chain A | A:13-141: C2 domain-like<br>A:142-721: FabD/lysophospholipase-like | A:13-141: PLC-like (P variant)<br>A:142-721: Lysophospholipase | 24.24% with 2mh1 |
| Synaptotagmin-1, C2A domain | 2k45 | C2 domain-like | Synaptotagmin-like (S variant) | 36.92% with 4v11 |
| Protein kinase C alpha, C2 domain | 4dnl | C2 domain-like | Synaptotagmin-like (S variant) | 33.08% with 2k45 |
| Synaptotagmin-1, C2B domain | 4v11 | C2 domain-like | Synaptotagmin-like (S variant) | 36.92% with 2k45 |
| Protein kinase C epsilon, C2 domain | 1gmi | C2 domain-like | PLC-like (P variant) | 23.38% with 4v11 |
| Neutrophil cytosol factor 4 (p40phox) | 1h6h | PX domain | PX domain | 25.68% with 1kq6 |
| Neutrophil cytosol factor 1 (p47phox) | 1kq6 | PX domain | PX domain | 25.68% with 1h6h |
| Early endosome antigen 1 FYVE domain | 1joc | A:1289-1347: Left-handed parallel coiled-coil<br>A:1348-1411: FYVE/PHD zinc finger | A:1289-1347: Eea1 homodimerisation domain<br>A:1348-1411: FYVE, a phosphatidylinositol-3-phosphate binding domain | 40.3% with 1vfy |
| Vps27 FYVE domain | 1vfy | FYVE/PHD zinc finger | FYVE, a phosphatidylinositol-3-phosphate binding domain | 43.28% with 1dvp |
| C1 domain of protein kinase C delta | 3uej, chain A | Cysteine-rich domain | C1 domain | 49.23% with 1tbn |
| Epsin ENTH domain | 1h0a | alpha-alpha superhelix | ENTH domain | 18.92% with 1la4 |
| C2 domain of coagulation factor V | 1czs | Galactose-binding domain-like | Discoidin domain (fa58c, coagulation factor 5/8 c-terminal domain) | 89.38% with 1sdd |
| C2 domains of coagulation factor Va | 1sdd | A:1-296: Cupredoxin-like<br>A:181-296: Cupredoxin-like<br>B:1863-2024: Galactose-binding domain-like<br>B:2025-2182: Galactose-binding domain-like | A:1-180: Multidomain cupredoxins<br>A:181-296: Multidomain cupredoxins<br>B:1863-2024: Discoidin domain (fa58c, coagulation factor 5/8 c-terminal domain)<br>B:2025-2182: Discoidin domain (fa58c, coagulation factor 5/8 c-terminal domain) | 89.38% with 1czs |
| Coagulation factor VIII | 2r7e | A:1-725: No SCOP2 classification is available<br>B:1689-2170: No SCOP2 classification is available<br>B:2171-2329: Galactose-binding domain-like | A:1-725: No SCOP2 classification is available<br>B:1689-2170: No SCOP2 classification is available<br>B:2171-2329: Discoidin domain (fa58c, coagulation factor 5/8 c-terminal domain) | 42.33% with 1czs |
| Annexin V | 1anx, chain A | Annexin | Annexin | 32.49% with 1dk5 |
| Equinatoxin II | 1iaz, chain A | Cytolysin/lectin | Anemone pore-forming cytolysin | 67.43% with 1gwy |

| | | | | |
|---|---|---|---|---|
| Prostaglandin H2 synthase 1 | 2ayl | Heme-dependent peroxidases | Myeloperoxidase-like | 28.33% with 1ffj |
| Pancreatic phospholipase A2, group IB | 1hn4, chain A | Phospholipase A2, PLA2 | Vertebrate phospholipase A2 | 53.23% with 1poa |
| Bee venom phospholipase A2 | 1poc | Phospholipase A2, PLA2 | Insect phospholipase A2 | 23.65% with 1joc |
| Phospholipase A2, group IIA | 5g3n, chain A | Phospholipase A2, PLA2 | Vertebrate phospholipase A2 | 50% with 4hg9 |
| Acidic phospholipase A2 1 | 1poa | Phospholipase A2, PLA2 | Vertebrate phospholipase A2 | 53.23% with 1hn4 |
| Snake phospholipase A2, group II | 1vap, chain A | Phospholipase A2, PLA2 | Vertebrate phospholipase A2 | 68.29% with 4hg9 |
| Snake phospholipase A2, group II, B | 4hg9, chain A | Phospholipase A2, PLA2 | Vertebrate phospholipase A2 | 68.29% with 1vap |
| Phosphatidylinositol-specific phospholipase C | 2ptd | TIM beta/alpha-barrel | Bacterial PLC | 19.05% with 1eaz |
| α-toxin (bacterial phospholipase C) | 1gyg, chain A | A:1-249: Phospholipase C/P1 nuclease<br>A:250-370: Lipase/lipooxygenase domain (plat/lh2 domain) | A:1-249: Phospholipase C<br>A:250-370: alpha-toxin, C-terminal domain | 24.24% with 1la4 |
| Arachidonate 15-lipoxygenase | 2p0m, chain A | A:2-112: Lipase/lipooxygenase domain (plat/lh2 domain)<br>A:113-663: Lipoxygenase | A:2-112: Lipoxygenase N-terminal domain-like<br>A:113-663: Lipoxygenase catalytic domain-like | 31.15% with 2fnq |
| 8R-Lipoxygenase | 2fnq, chain A | No SCOP2 classification is available | No SCOP2 classification is available | 31.15% with 2p0m |
| Signal peptidase I | 3iiq, chain A | SH3-like barrel | Type 1 signal peptidase | 27.27% with 2mh1 |
| Perfringolysin | 1pfo | A:103-368: MAC/CDC-like<br>A:390-500: Prealbumin-like | A:30-500: Perfringolysin-like | 19.05% with 4v11 |
| Hepatocyte growth factor-regulated tyrosine kinase substrate (Hrs) | 1dvp | A:1-145: alpha-alpha superhelix<br>A:149-219: FYVE/PHD zinc finger | A:1-145: VHS domain<br>A:149-219: FYVE, a phosphatidylinositol-3-phosphate binding domain | 43.28% with 1vfy |
| Prothrombin, GLA domain | 1nl1 | A:1-65: GLA-domain<br>A:66-147: Kringle-like | A:1-65: GLA-domain<br>A:66-147: Kringle modules | 27.03% with 1tbn |
| Sorting nexin-3 | 5f0p | A:12-469: No SCOP2 classification is available<br>B:8-300: Immunoglobulin-like beta-sandwich<br>C:4-150: PX domain<br>D:551-565: No SCOP2 classification is available | A:12-469: No SCOP2 classification is available<br>B:8-300: Vps26-like<br>C:4-150: PX domain<br>D:551-565: No SCOP2 classification is available | 23.53% with 1s6x |
| Beta-2-glycoprotein 1 | 1c1z | Sushi/Complement control module/SCR domain | Sushi/CCP/SCR domain | 19.64% with 1faq |
| Alpha-tocopherol transfer protein | 1oiz, chain A | A:9-90: UBA-type 3-helical bundle<br>A:91-274: SpoIIaa-like | A:9-90: CRAL/TRIO N-terminal domain<br>A:91-274: CRAL/TRIO domain | 18.18% with 1la4 |
| Annexin 24 | 1dk5, chain A | Annexin | Annexin | 32.49% with 1anx |
| Protein kinase C gamma type C1B domain | 1tbn | Cysteine-rich domain | C1 domain | 49.23% with 3uej |
| Protein kinase C gamma type C1A domain | 2e73 | Cysteine-rich domain | C1 domain | 38.46% with 3uej |
| RAF-1 proto-oncogene serine/threonine-protein | 1faq | Cysteine-rich domain | C1 domain | 28.57% with 3uej |

| | | | | |
|---|---|---|---|---|
| kinase | | | | |
| Eosinophil cationic protein | 4x08, chain A | RNase A-like | Ribonuclease A-like | 22.81% with 1faq |
| Antimicrobial peptide kalata B1 | 2mh1 | Knottins (small inhibitors, toxins, lectins) | Kalata B1 | 27.27% with 3iiq |
| Kappa-theraphotoxin-Scg1a | 1la4 | Knottins (small inhibitors, toxins, lectins) | Spider toxins | 36.36% with 1s6x |
| Dual adapter for phosphotyrosine and 3-phosphotyrosine and 3-phosphoinositide | 1fao | PH domain-like beta(6)-barrel | Pleckstrin-homology domain (PH domain) | 36.54% with 1eaz |
| Pleckstrin homology domain-containing family A member 1 | 1eaz | PH domain-like beta(6)-barrel | Pleckstrin-homology domain (PH domain) | 36.54% with 1fao |
| Sticholysin II | 1gwy, chain A | Cytolysin/lectin | Anemone pore-forming cytolysin | 93.14% with 2ks4 |
| Sticholysin I | 2ks4 | Cytolysin/lectin | Anemone pore-forming cytolysin | 93.14% with 1gwy |
| Matrix protein VP40 | 1es6 | EV matrix protein-like | EV matrix protein | 18.79% with 1kq6 |
| | | **Test set 1** | | |
| Retinoid isomerohydrolase | 3fsn | 7-bladed beta-propeller | Retinoid isomerase RPE65-like | 22.35% with 3npe |
| Voltage sensor toxin VSTx1 | 1s6x | No SCOP2 classification is available | No SCOP2 classification is available | 36.36% with 1la4 |
| Cytotoxin 2 | 1ffj | Snake toxin-like | Snake venom toxins | 28.33% with 2ayl |
| Sphingomyelinase C | 2ddr, chain A | DNase I-like | Sphingomyelin phosphodiesterase-like | 18.48% with 1jss |
| Glycolipid transfer protein | 3rzn | Glycolipid transfer protein, GLTP | Glycolipid transfer protein, GLTP | 18.33% with 1ffj |
| Cholesterol-regulated Start protein 4 | 1jss, chain A | TBP-like | STAR domain | 18.48% with 2ddr |
| Ceramide transfer protein, PH domain | 2rsg | PH domain-like beta(6)-barrel | Pleckstrin-homology domain (PH domain) | 25% with 1eaz |
| Phosphatidylinositol transfer protein beta isoform | 2a1l | TBP-like | Phoshatidylinositol transfer protein, PITP | 22.86% with 3uej |
| | | **Test set 2** | | |
| Cholesterol oxidase | 1coy | FAD/NAD(P)-binding domain | GMC oxidoreductase-like | 21.21% with 2mh1 |
| Cytochrome P450 3A4 | 1tqn | Cytochrome P450 | Cytochrome P450 | 18.63% with 1fao |
| 9-cis-epoxycarotenoid dioxygenase 1, chloroplastic | 3npe | 7-bladed beta-propeller | Retinoid isomerase RPE65-like | 27.27% with 2mh1 |
| Monoglyceride lipase MGLL | 3jw8, chain A | No SCOP2 classification is available | No SCOP2 classification is available | 23.19% with 3uej |
| Dihydroorotate dehydrogenase | 3w7r | TIM beta/alpha-barrel | FMN-linked oxidoreductases | 20.59% with 1s6x |
| Phosphatase PTEN | 5bzz, chain A | A:14-187: (Phosphotyrosine protein) phosphatases II<br>A:188-351: C2 domain-like | A:14-187: Dual specificity phosphatase-like<br>A:188-351: PLC-like (P variant) | 21.21% with 2mh1 |
| (S)-mandelate dehydrogenase | 6bfg | TIM beta/alpha-barrel | FMN-linked oxidoreductases | 21.21% with 2mh1 |
| (S)-mandelate dehydrogenase, homotetramer | Homotetramer biological assembly of 6bfg | TIM beta/alpha-barrel | FMN-linked oxidoreductases | 21.21% with 2mh1 |
| L-amino acid deaminase | 5hxw, chain A | No SCOP2 classification is available | No SCOP2 classification is available | 16.42% with 1vfy |

| | | | | |
|---|---|---|---|---|
| Intestinal fatty acid binding protein | 3akm, chain A | Lipocalins | Fatty acid binding protein-like | 19.71% with 3iiq |
| Phosphatidylinositol 4,5-bisphosphate 3-kinase | 4ovu | A:1-143: No SCOP2 classification is available A:144-322: beta-Grasp<br>A:357-522: C2 domain-like<br>A:525-725: alpha-alpha superhelix<br>A:726-1092: Protein kinase-like (PK-like)<br>B:327-433: SH2-like<br>B:434-598: Left-handed antiparallel coiled-coil | A:1-143: No SCOP2 classification is available<br>A:144-322: Ras-binding domain, RBD<br>A:357-522: PLC-like (P variant)<br>A:525-725: Phoshoinositide 3-kinase (PI3K) helical domain<br>A:726-1092: Phoshoinositide 3-kinase (PI3K), catalytic domain<br>B:327-433: SH2 domain<br>B:434-598: PI3K inter-SH2 coiled-coil domain | 22.22% with 1faq |
| Phosphatidylcholine transfer protein | 1ln1 | TBP-like | STAR domain | 22.86% with 2mh1 |

**Table S6.** The hyper-parameters that were sampled as these are named and described in scikit-learn [23], LightGBM [24], and XGBoost [110] packages and their ranges that were searched in randomized and grid search cross-validation and the final best hyper-parameters for each machine learning classifier.

| Classifier | Hyper-parameter ranges selected herein | Best hyper-parameters identified by hyper-parameter optimization |
|---|---|---|
| Decision tree | 'criterion': ['gini', 'entropy'], 'max_features': [3-150, 'sqrt', 'auto', 'log2', None], 'splitter': ['best', 'random'], 'max_depth': [None, 3-15], 'min_samples_split': [2-20, 0.1], 'min_impurity_decrease': [0-0.001], 'min_samples_leaf': [1-20, 0.1], 'class_weight' : ['balanced'] | 'class_weight': 'balanced', 'criterion': 'gini', 'max_depth': 11, 'max_features': 121, 'min_impurity_decrease': 3e-05, 'min_samples_leaf': 1, 'min_samples_split': 5, 'splitter': 'best' |
| Extreme gradient boosting | 'max_depth': [1-10], 'min_child_weight': [1-20], 'gamma': [0-0.5], 'subsample': [0.6-1], 'colsample_bynode': [0.6-1], 'colsample_bylevel': [0.6-1], 'colsample_bytree': [0.6-1], 'reg_alpha': [0-100], 'reg_lambda': [0-100], 'grow_policy': [depthwise, lossguide], 'learning_rate': [0.001-0.5], 'n_estimators': [50-1000], 'max_delta_step': [0-10], 'sampling_method': ['uniform'], 'scale_pos_weight': [17.024], 'tree_method': ['gpu_hist'] | 'colsample_bylevel': 0.96, 'colsample_bynode': 0.74, 'colsample_bytree': 0.84, 'gamma': 0, 'grow_policy': 'depthwise', 'learning_rate': 0.1, 'max_delta_step': 1.6, 'max_depth': 4, 'min_child_weight': 1, 'n_estimators': 160, 'reg_alpha': 0.1, 'reg_lambda': 16, 'sampling_method': 'uniform', 'scale_pos_weight': 17.024, 'subsample': 0.83, 'tree_method': 'gpu_hist' |
| Extremely randomized trees | 'criterion': ['gini', 'entropy'], 'max_features': [3-50, 'auto', 'log2', None], 'n_estimators': [50-1000], 'max_depth': [None, 3-50], 'min_samples_split': [2-40, 0.1], 'min_impurity_decrease': [0-0.001], 'warm_start': [True, False], 'min_samples_leaf': [1-20, 0.1], 'class_weight' : ['balanced'], 'bootstrap': [True, False] | 'bootstrap': False, 'class_weight': 'balanced', 'criterion': 'entropy', 'max_depth': 17, 'max_features': None, 'min_impurity_decrease': 9.5e-09, 'min_samples_leaf': 6, 'min_samples_split': 15, 'n_estimators': 86, 'warm_start': False |
| Gaussian naive Bayes | 'var_smoothing': [1e-11-0.1] | 'var_smoothing': 1e-11 |
| Gaussian processes | 'warm_start': [True, False], 'kernel': [RBF(), ConstantKernel(), DotProduct(),WhiteKernel(), RationalQuadratic(), ExpSineSquared(), Exponentiation(), Matern(), PairwiseKernel()], 'max_iter_predict': [50-300], 'n_restarts_optimizer': [1-3] | 'kernel': RBF(), 'max_iter_predict': 300, 'n_restarts_optimizer': 3, 'warm_start': True |
| Gradient boosting | 'loss': ['deviance', 'exponential'], 'learning_rate': [0.0001-1], 'n_estimators': [50-1000], 'subsample': [0.5-1], 'criterion': ['friedman_mse', 'mse', 'mae'], 'min_samples_split': [2-20, 0.1], 'min_samples_leaf': [1-20, 0.1, 0.2], 'max_depth': [None, 3-15], 'min_impurity_decrease': [0-0.001], 'max_features': [3-150, 'auto', 'log2', None], 'warm_start': [True, False], 'tol': [1e-8-0.1] | 'warm_start': False, 'tol': 1e-07, 'subsample': 0.85, 'n_estimators': 900, 'min_samples_split': 2, 'min_samples_leaf': 0.1, 'min_impurity_decrease': 0, 'max_features': 'auto', 'max_depth': 5, 'loss': 'deviance', 'learning_rate': 0.3, 'criterion': 'mse' |
| k-nearest neighbors | 'n_neighbors': [1-31], 'algorithm': ['ball_tree', 'kd_tree', 'brute'], 'leaf_size': [1-100], 'metric': ['cityblock', 'euclidean', 'l1', 'l2', 'manhattan'], 'weights': ['uniform', 'distance'], 'p': [1-3] | 'algorithm': 'ball_tree', 'leaf_size': 1, 'metric': 'cityblock', 'n_neighbors': 1, 'p': 1, 'weights': 'uniform' |
| Light gradient boosting machine | 'n_estimators': [10-1000], 'min_child_weight': [1e-5-0], 'min_child_samples': [2-200], 'subsample': [0.6-1], 'colsample_bytree': [0.6-1], 'subsample_freq': [5], 'num_leaves': [3-300], 'learning_rate': [0.001-0.5], 'reg_alpha': [0-100], 'reg_lambda': [0- 100], 'metric': ['binary_logloss'], 'scale_pos_weight': [17.024], 'objective': ['binary'] | 'subsample_freq': 5, 'subsample': 1.0, 'scale_pos_weight': 17.024, 'reg_lambda': 0, 'reg_alpha': 5, 'objective': 'binary', 'num_leaves': 58, 'n_estimators': 820, 'min_child_weight': 0.0001, 'min_child_samples': 96, 'metric': 'binary_logloss', 'learning_rate': 0.1174, 'colsample_bytree': 0.8 |
| Linear discriminant analysis | 'solver': ['svd', 'lsqr', 'eigen'], 'shrinkage': [1e-5-0.9], 'tol' : [1e-11-0.1] | 'solver': 'svd', 'tol': 0.1 |
| Linear support vector machine | 'penalty': ['l1', 'l2'], 'loss': ['hinge', 'squared_hinge'], 'dual': [True, False], 'C': [0.001-10], 'fit_intercept': [True, False], 'max_iter': [500-3000], 'tol': [1e-9-0.1], 'class_weight': ['balanced'] | 'C': 0.0052, 'class_weight': 'balanced', 'dual': False, 'fit_intercept': True, 'loss': 'squared_hinge', 'max_iter': 580, 'penalty': 'l1', 'tol': 1.5e-07 |
| Logistic regression | 'solver': ['liblinear', 'newton-cg', 'lbfgs', 'sag', 'saga'], 'C': [0.001-10], 'penalty': ['l1', 'l2', 'elasticnet', 'none'], 'l1_ratio': [0.1- 0.9], 'max_iter': [500-3000], 'tol': [1e-10-0.1], 'class_weight': ['balanced'] | 'C': 0.0086, 'class_weight': 'balanced', 'l1_ratio': 0.21, 'max_iter': 2200, 'penalty': 'elasticnet', 'solver': 'saga', 'tol': 1e-09 |
| Multilayer | 'hidden_layer_sizes': (10-150, 10-150, 5-150), 'activation': | 'activation': 'tanh', 'alpha': 0.5, 'batch_size': 'auto', |

| | | |
|---|---|---|
| perceptron | ['identity', 'logistic', 'tanh', 'relu'], 'alpha': [1e-5-10], 'learning_rate': ['constant', 'invscaling', 'adaptive'], 'tol': [1e-6-0.1], warm_start': [True, False], 'batch_size': ['auto', 50- 2000], 'solver': ['lbfgs', 'sgd', 'adam'], 'max_iter': [100-600], 'early_stopping': [True, False], 'epsilon': [1e-9-1e-5] | 'early_stopping': False, 'epsilon': 1e-07, 'hidden_layer_sizes': (35, 150, 5), 'learning_rate': 'invscaling', 'max_iter': 190, 'solver': 'adam', 'tol': 1e-05, 'warm_start': True |
| Nearest centroid | 'metric': ['cityblock', 'euclidean', 'l1', 'l2', 'manhattan'], 'shrink_threshold': [1e-5-2, None] | 'metric': 'cityblock', 'shrink_threshold': 0.045 |
| v-support vector machine | 'nu': [1e-5-0.1], 'kernel': ['linear', 'poly', 'rbf', 'sigmoid'], 'degree': [1-8], 'gamma': [1e-5-1, 'auto', 'scale'], 'coef0': [0-10], 'shrinking': [True, False], 'tol': [1e-6-0.1], 'max_iter': [10000], 'class_weight': ['balanced'] | 'class_weight': 'balanced', 'coef0': 2.4, 'degree': 4, 'gamma': 0.0001, 'kernel': 'poly', 'max_iter': 10000, 'nu': 0.055, 'shrinking': True, 'tol': 0.45 |
| Passive-aggressive | 'C': [1e-5-10], 'fit_intercept': [True, False],'max_iter': [500-3000], 'warm_start': [True, False], 'tol': [1e-5-0.1], 'loss': ['hinge', 'squared_hinge'], 'class_weight' : ['balanced'] | 'C': 6e-05, 'class_weight': 'balanced', 'fit_intercept': True, 'loss': 'squared_hinge', 'max_iter': 2600, 'tol': 5.5e-05, 'warm_start': True |
| Perceptron | 'penalty': ['l2', 'l1', 'elasticnet'], 'alpha': [1e-4-0.1], 'max_iter': [30, 3000], 'eta0': [1e-9-1], 'warm_start': [True, False], 'tol': [1e-11- 0.1], 'class_weight' : ['balanced']} | 'alpha': 0.008, 'class_weight': 'balanced', 'eta0': 9e-05, 'max_iter': 40, 'penalty': 'l2', 'tol': 1e-11, 'warm_start': True |
| Quadratic discriminant analysis | 'reg_param': [1e-10-0.1], 'tol' : [1e-12-0.1] | 'reg_param': 0.00105, 'tol': 1e-09 |
| Radius nearest neighbors | 'radius': [100-10000],'weights': ['uniform', 'distance'], 'algorithm': ['ball_tree', 'kd_tree', 'brute'], 'leaf_size': [1-100], 'metric': ['cityblock', 'euclidean', 'l1', 'l2', 'manhattan'], 'p': [1-3], 'outlier_label': ['most_frequent'] | 'algorithm': 'brute', 'leaf_size': 1, 'metric': 'l1', 'outlier_label': 'most_frequent', 'p': 1, 'radius': 234, 'weights': 'distance' |
| Random forest | 'criterion': ['gini', 'entropy'], 'max_features': [3-50, 'auto', 'log2', None], 'n_estimators': [50-1000], 'max_depth': [None, 3-40], 'min_samples_split': [2-20, 0.1], 'min_impurity_decrease': [0-0.001], 'warm_start': [True, False], 'min_samples_leaf': [1-30, 0.1], 'class_weight' : ['balanced'], 'bootstrap': [True, False] | 'bootstrap': True, 'class_weight': 'balanced', 'criterion': 'entropy', 'max_depth': 38, 'max_features': None, 'min_impurity_decrease': 0, 'min_samples_leaf': 9, 'min_samples_split': 2, 'n_estimators': 490, 'warm_start': True |
| Stochastic gradient descent | 'loss': ['hinge', 'log', 'modified_huber', 'squared_hinge', 'perceptron', 'squared_loss', 'huber', 'epsilon_insensitive', 'squared_epsilon_insensitive'], 'alpha': [1e-4-100], 'penalty': ['l1', 'l2', None, 'elasticnet'], 'max_iter': [500-3000], 'warm_start': [True, False], 'tol': [1e-6-0.9], 'l1_ratio': [0.1- 0.9], 'class_weight': ['balanced'] | 'alpha': 0.01, 'class_weight': 'balanced', 'l1_ratio': 0.5, 'loss': 'perceptron', 'max_iter': 2600, 'penalty': 'elasticnet', 'tol': 0.1, 'warm_start': False |
| Support vector machine | 'C': [0.001-10], 'kernel': ['linear', 'poly', 'rbf', 'sigmoid'], 'degree': [1- 8], 'gamma': [0.001-10, 'auto', 'scale'], 'coef0': [0-10], 'shrinking': [True, False], 'tol': [1e-11-0.1], 'max_iter': [10000], 'class_weight': ['balanced'] | 'C': 1, 'class_weight': 'balanced', 'coef0': 1, 'degree': 2, 'gamma': 'scale', 'kernel': 'poly', 'max_iter': 10000, 'shrinking': True, 'tol': 1e-06 |

**Table S7.** The test set predictions of the 21 classifiers and the chosen voting classifier for the first test set after data selection. The best score for each metric is highlighted with bold.

| Classifier | TN | FN | TP | FP | Precision macro | Recall macro | $F_1$ score macro | $F_2$ score macro | PR AUC | g-mean macro | MCC |
|---|---|---|---|---|---|---|---|---|---|---|---|
| Decision tree | 463 | 26 | 18 | 10 | 0.79 | 0.69 | 0.73 | 0.71 | 0.55 | 0.69 | 0.48 |
| Extreme gradient boosting | 464 | 20 | 24 | 9 | 0.84 | 0.76 | 0.80 | 0.78 | 0.66 | 0.76 | 0.60 |
| Extremely randomized trees | 462 | 22 | 22 | 11 | 0.81 | 0.74 | 0.77 | 0.75 | 0.60 | 0.74 | 0.54 |
| Gaussian naive Bayes | 437 | 16 | 28 | 36 | 0.70 | 0.78 | 0.73 | 0.76 | 0.55 | 0.78 | 0.47 |
| Gaussian processes | 468 | 35 | 9 | 5 | 0.79 | 0.60 | 0.63 | 0.61 | 0.46 | 0.60 | 0.33 |
| Gradient boosting | 468 | 27 | 17 | 5 | **0.86** | 0.69 | 0.74 | 0.70 | 0.61 | 0.69 | 0.52 |
| k-nearest neighbors | 458 | 27 | 17 | 15 | 0.74 | 0.68 | 0.70 | 0.69 | 0.48 | 0.68 | 0.41 |
| Light gradient boosting machine | 463 | 16 | 28 | 10 | 0.85 | 0.81 | 0.83 | 0.82 | 0.70 | 0.81 | 0.66 |
| Linear discriminant analysis | 456 | 19 | 25 | 17 | 0.78 | 0.77 | 0.77 | 0.77 | 0.60 | 0.77 | 0.54 |
| Linear support vector machine | 445 | 3 | 41 | 28 | 0.79 | **0.94** | 0.85 | **0.89** | **0.77** | **0.94** | **0.72** |
| Logistic regression | 452 | 6 | 38 | 21 | 0.82 | 0.91 | 0.85 | **0.89** | 0.76 | 0.91 | **0.72** |
| Multilayer perceptron | 435 | 18 | 26 | 38 | 0.68 | 0.76 | 0.71 | 0.73 | 0.52 | 0.76 | 0.43 |
| Nearest centroid | 444 | 27 | 17 | 29 | 0.66 | 0.66 | 0.66 | 0.66 | 0.40 | 0.66 | 0.32 |
| ν-support vector machine | 397 | 29 | 15 | 76 | 0.55 | 0.59 | 0.55 | 0.57 | 0.28 | 0.59 | 0.13 |
| Passive-aggressive | 385 | 22 | 22 | 88 | 0.57 | 0.66 | 0.58 | 0.61 | 0.37 | 0.66 | 0.21 |
| Perceptron | 376 | 22 | 22 | 97 | 0.56 | 0.65 | 0.57 | 0.60 | 0.36 | 0.65 | 0.20 |
| Quadratic discriminant analysis | 464 | 32 | 12 | 9 | 0.75 | 0.63 | 0.66 | 0.64 | 0.45 | 0.63 | 0.36 |
| Radius nearest neighbors | 473 | 44 | 0 | 0 | 0.46 | 0.50 | 0.48 | 0.49 | 0.54 | 0.50 | 0.00 |
| Random forest | 462 | 26 | 18 | 11 | 0.78 | 0.69 | 0.73 | 0.70 | 0.54 | 0.69 | 0.47 |
| Stochastic gradient descent | 453 | 18 | 26 | 20 | 0.76 | 0.77 | 0.77 | 0.77 | 0.60 | 0.77 | 0.54 |
| Support vector machine | 463 | 29 | 15 | 10 | 0.77 | 0.66 | 0.70 | 0.67 | 0.50 | 0.66 | 0.42 |
| Voting ensembles | 459 | 10 | 34 | 14 | 0.84 | 0.87 | **0.86** | 0.87 | 0.75 | 0.87 | 0.71 |

Note that because not every sample in our dataset is experimentally tested, many false positives may be actually membrane-penetrating amino acids, resulting in lower score values.

**Table S8.** Peripheral membrane proteins of the test set, their PDB code, their experimentally known hydrophobic membrane-penetrating amino acids after data selection, and the true positive and false positive predictions of our ensemble classifier. Amino acid numbering is the same as in the PDB structures. With † we note the amino acids that were discarded because their amino acid depth value was more than 2.5 Å. With * we note the false positive predictions that are on the protein-membrane interface, indicating that are true positives in reality.

| Protein | PDB | Membrane-penetrating amino acids | True positives | False positives |
|---|---|---|---|---|
| Retinoid isomerohydrolase | 3fsn | F196, F200, I202†, F264, L265, W268, L270, W271 | Chain A: F200, L265, W268, L270, W271<br>Chain B: F196, F200, L265, W268, L270, W271 | Chain A: F262*<br>Chain B: F262* |
| Voltage sensor toxin VSTx1 | 1s6x | F5, M6, W7, W27, V29†, L30 | M6, W7 | W25, P33 |
| Cytotoxin 2 | 1ffj | L6, V7, P8, L9, F10, Y22†, M24, F25, M26, V27, P30, V32, P33, V34, I39, L47, L48, V49 | L6, V7, L9, F10, F25, M26, V27, P30, V32, V34, L47, L48, V49 | V41 |
| Sphingomyelinase C | 2ddr, chain A | W284, F285 | W284, F285 | |
| Glycolipid transfer protein | 3rzn | W142 | W142 | Y81, I143*, Y153* |
| Cholesterol-regulated Start protein 4 | 1jss, chain A | L124 | L124 | W91, M196* |
| Ceramide transfer protein, PH domain | 2rsg | W33, Y36 | W33, Y36 | I37*, W40*, F81* |
| Phosphatidylinositol transfer protein beta isoform | 2a1l | W202, W203 | W202, W203 | M74* |

**Table S9.** The peripheral membrane proteins of the test set, their PDB code, their experimentally known hydrophobic membrane-penetrating amino acids after data selection, the predictions of our ensemble classifier keeping all amino acid types, the PPM predictions, and the MODA predictions. Amino acid numbering is the same as in the PDB structures.

| Protein | PDB | Membrane-penetrating amino acids | Our classifier | PPM | MODA |
|---|---|---|---|---|---|
| Retinoid isomerohydrolase | 3fsn | F196, F200, I202, F264, L265, W268, L270, W271 | Chain A: F200, F262, L265, W268, L270, W271<br>Chain B: F196, F200, F262, L265, W268, L270, W271 | Chain A: -<br>Chain B: P109, F196, N199, F200, F262, L265, S266, W268, S269, L270, W271 | Chain A: R33, S54, E55, V99, A107, F108, F196, G197, N199, F200, S201, R234, F235, L261, F262, L265, S266, S267, W268, S269, L270, W271, G272<br>Chain B: R33, F108, P109, F196, G197, K198, N199, F200, R234, F235, L261, F262, L265, S266, S267, W268, S269, L270, W271, G272 |
| Voltage sensor toxin VSTx1 | 1s6x | F5, M6, W7, W27, V29, L30 | E1, M6, W7, K8, D18, R24, W25, K26, S32, P33 | F5, M6, W25, W27 | K4, F5, M6, W7, V20, C21, S22, S23, R24, W25, K26, W27, V29, L30, S32, P33, F34 |
| Cytotoxin 2 | 1ffj | L6, V7, P8, L9, F10, Y22, M24, F25, M26, V27, P30, V32, P33, V34, I39, L47, L48, V49 | L6, V7, L9, F10, F25, M26, V27, A28, P30, H31, V32, V34, R36, V41, C42, K44, L47, L48, V49 | L6, V7, P8, L9, F25, M26, V27, A28, A29, P30, H31, V32, P33, V34, K35, S46, L47, L48, V49 | K4, K5, L6, V7, P8, L9, F10, S11, K12, Y22, M24, F25, M26, V27, A28, A29, P30, V32, P33, V34, K35, P43, K44, S45, S46, L47, L48, V49, K50, Y51 |
| Sphingomyelinase C | 2ddr, chain A | W284, F285 | W284, F285 | L128, W284, F285 | N23, L24, Y25, P26, N27, C123, L128, Y242, N243, F244, P245, T282, S283, W284, F285, Q286, K287, Y288 |
| Glycolipid transfer protein | 3rzn | W142 | Y81, W142, I143, K146, Y153 | P40, V41, T43, P44, W142, I143, I147 | G141, W142, I143, V144, Q145, K146, I147, Q149, A150, Y153, V209 |
| Cholesterol-regulated Start protein 4 | 1jss, chain A | L124 | W91, L124, M196 | L124, I126, P198, S200 | L124, N125, R222 |
| Ceramide transfer protein, PH domain | 2rsg | W33, Y36 | W33, N35, Y36, I37, H38, G39, W40, F81, E83 | Y36, I37, H38, G39, W40, F81 | S31, K32, W33, T34, N35, Y36, I37, H38, G39, W40, H79, D80, F81, D82, R85, R98 |
| Phosphatidylinositol transfer protein beta isoform | 2a1l | W202, W203 | M74, W202, W203 | M74, W203 | M74, I75, W202, W203, G204 |

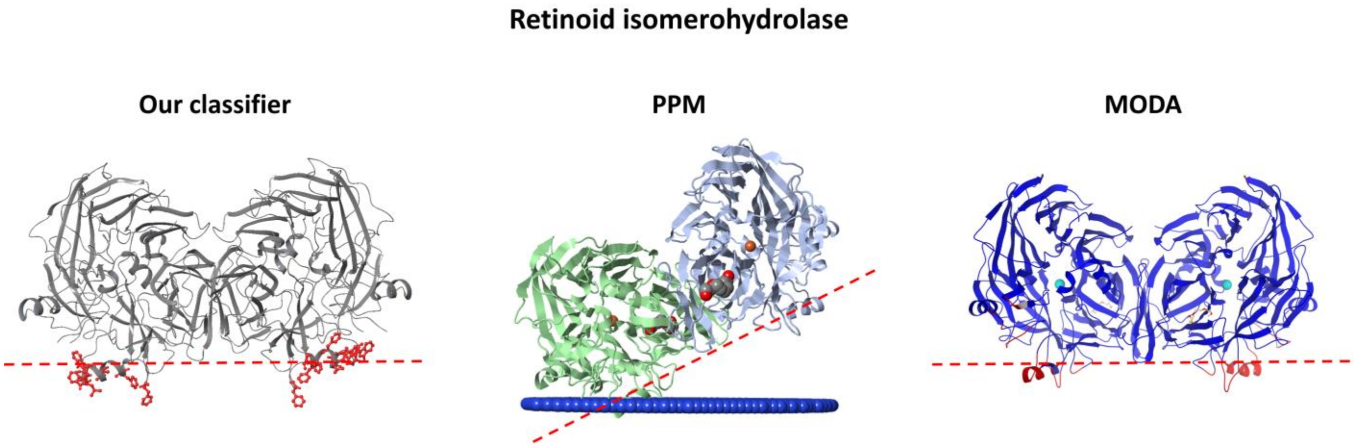


**Figure S3.** Comparison of the predictions provided from our classifier, PPM, and MODA for the retinoid isomerohydrolase homodimer. Both our classifier and MODA correctly found the protein-membrane regions in both chains, while PPM places only one chain in the membrane. The membrane is depicted with a red dotted line according to Ref. [96].

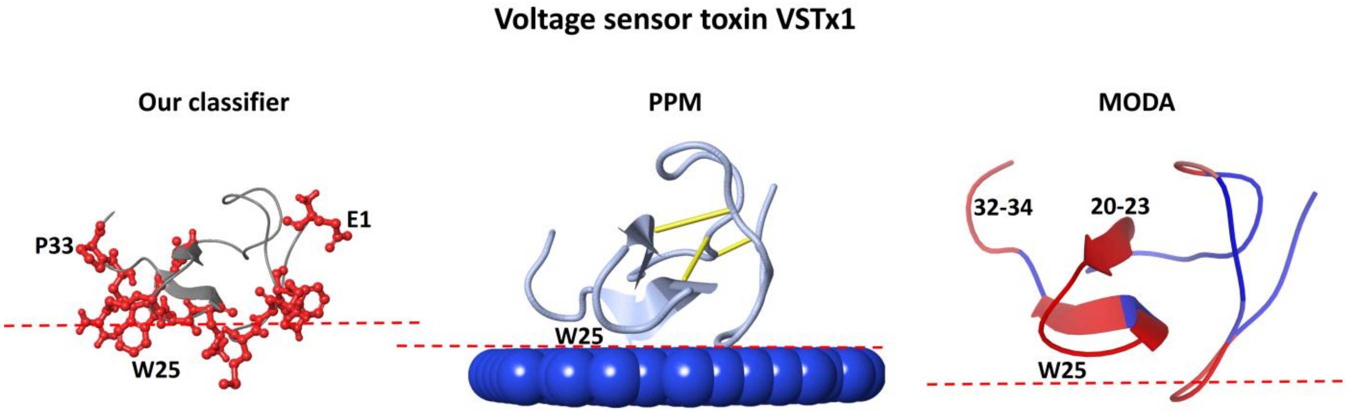


**Figure S4.** Comparison of the predictions provided from our classifier, PPM, and MODA for the VSTx1 toxin. Correct predictions for every tool, with every tool falsely predicting W25, our classifier falsely predicting the N- and C-terminal, and MODA falsely predicting the beta sheet on the opposite side of the protein-membrane interface and the C-terminal. The membrane is depicted with a red dotted line according to Ref. [97].

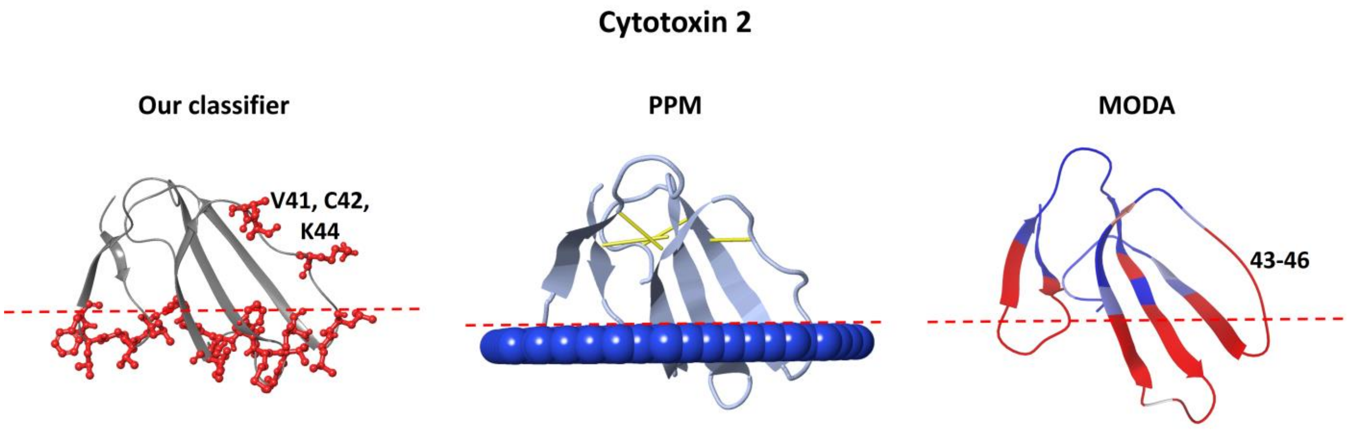


**Figure S5.** Comparison of the predictions provided from our classifier, PPM, and MODA for the cytotoxin 2. Correct predictions for every tool, with false positives for our classifier and MODA in the 41-46 region. The membrane is depicted with a red dotted line according to Ref. [100].

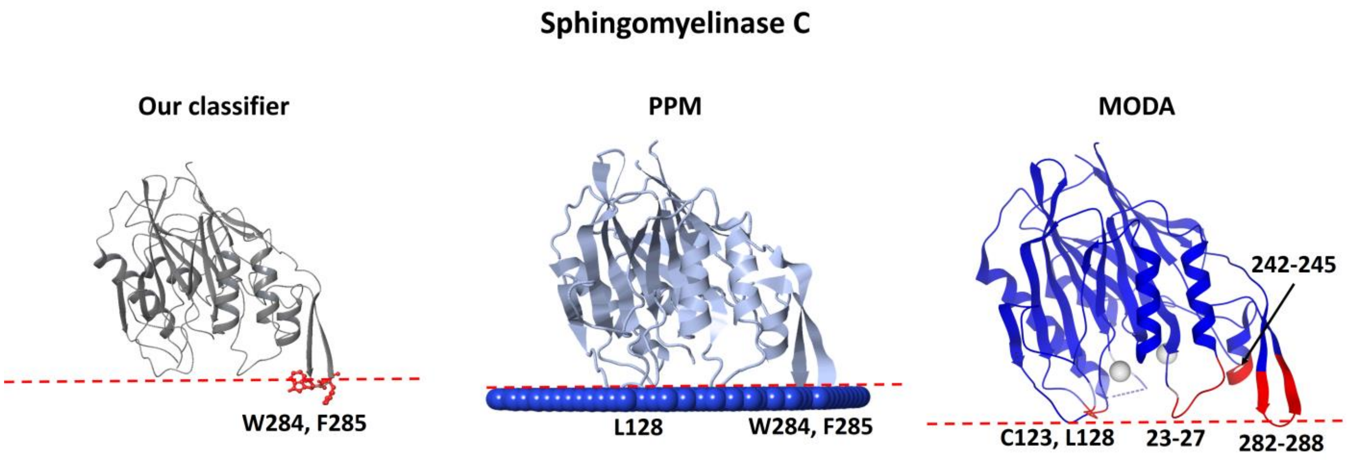


**Figure S6.** Comparison of the predictions provided from our classifier, PPM, and MODA for the sphingomyelinase C. Correct predictions for every tool, with PPM and MODA predicting the insertion of other loops which are aligned with the experimental amino acids. The membrane is depicted with a red dotted line according to Ref. [101].

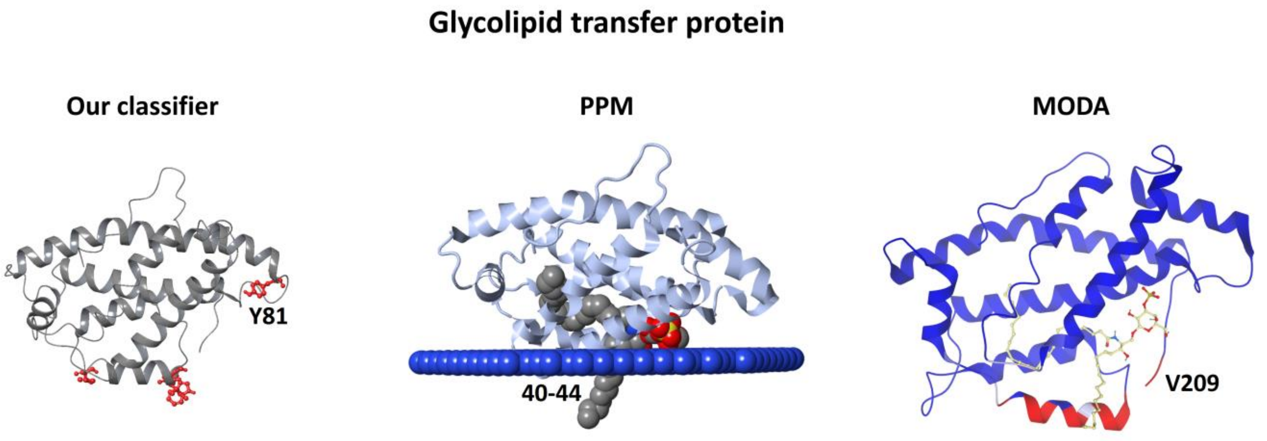


**Figure S7.** Comparison of the predictions provided from our classifier, PPM, and MODA for the glycolipid transfer protein. Correct predictions for every tool, with our classifier falsely predicting Y81, MODA falsely predicting the C-terminal, and PPM suggesting the insertion of the P40-P44 region.

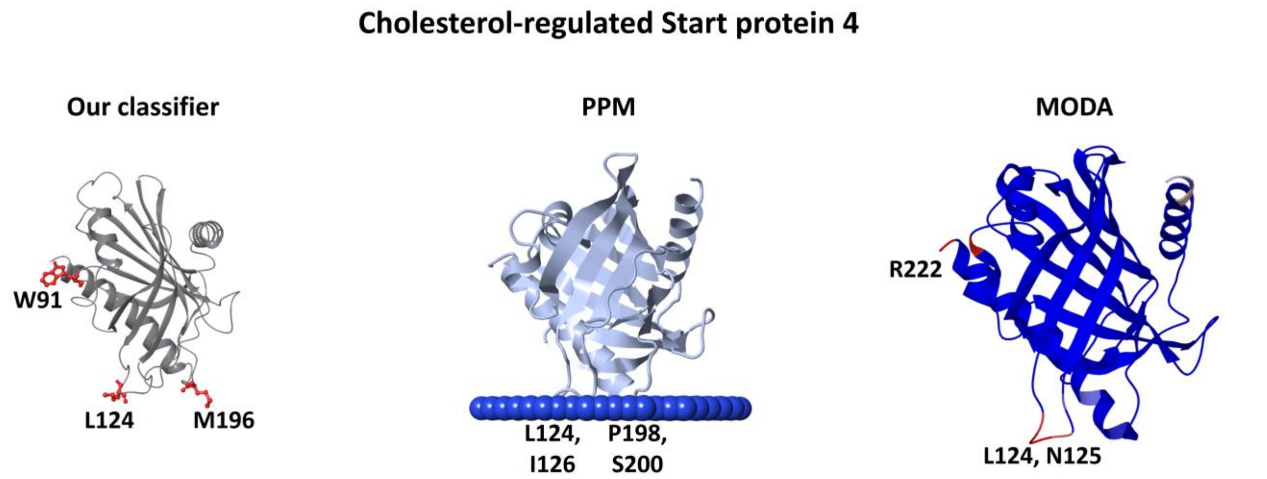


**Figure S8.** Comparison of the predictions provided from our classifier, PPM, and MODA for the cholesterol-regulated Start protein 4. All tools, predicted the experimentally proven membrane-penetrating amino acid L124, with our classifier and PPM additionally predicting the 196-200 region, MODA falsely predicting the C-terminus, and our classifier falsely predicting amino acid W91.

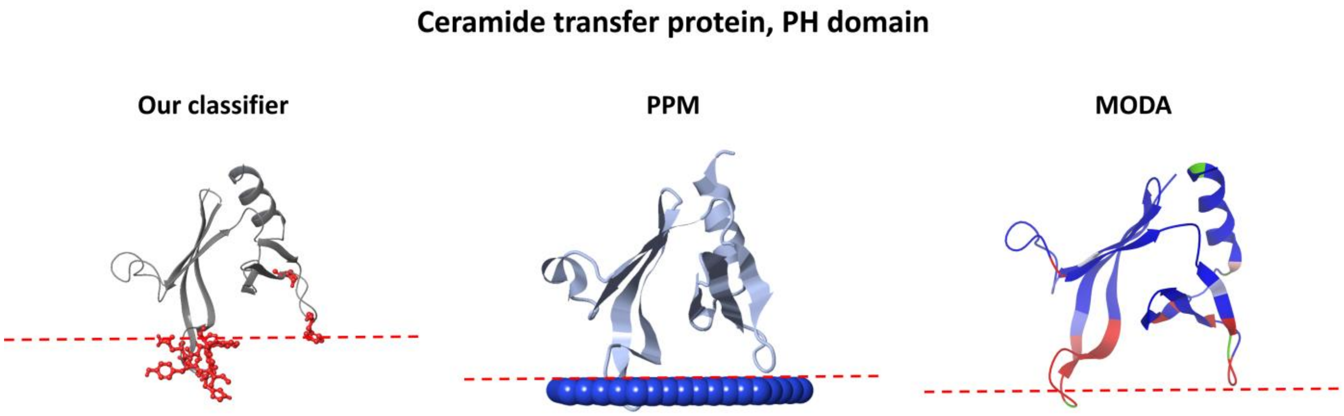


**Figure S9.** Comparison of the predictions provided from our classifier, PPM, and MODA for the ceramide transfer protein. Correct prediction for every tool. The membrane is depicted with a red dotted line according to Ref. [105].

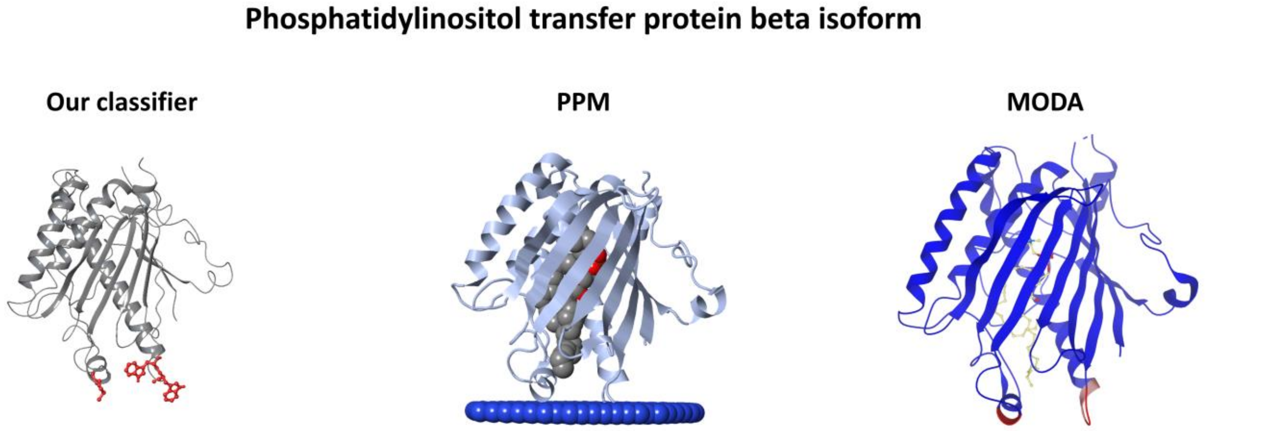


**Figure S10.** Comparison of the predictions provided from our classifier, PPM, and MODA for the phosphatidylinositol transfer protein beta isoform PH domain. Correct prediction for every tool.

**Table S10.** The peripheral membrane proteins of the test set 2, their PDB code, their experimentally known membrane-penetrating regions, the predictions of our ensemble classifier, the PPM predictions, the MODA predictions, and the experimental method used to determine experimentally known membrane-penetrating regions. Amino acid numbering is the same as in the PDB structures.

| **Protein** | **PDB** | **Membrane-penetrating regions** | **Our classifier** | **PPM** | **MODA** | **Methods & References** |
|---|---|---|---|---|---|---|
| Cholesterol oxidase | 1coy | Around amino acid M81 | M332, W333, Y436, Y437 | M81, H331, M332, W333, Y372, Y437 | R4, H331, M332, W333, G425, Y436, Y437 | FLq [111] |
| Cytochrome P450 3A4 | 1tqn | F-G loop and helices (218-238) and A-anchor (44-47) | F46, F228, I232, V235, L236 | H28, L44, P45, F46, L47, L51, P218, F219, S222, V225, F226, F228, L229, I232, L233, L236, I238 | H28, P43, L44, P45, F46, L47, L51, P218, S222, V225, F226, P227, F228, L229, P231, I232, L233, V235, L236, N237, I238 | LD [112] |
| 9-cis-epoxycarotenoid dioxygenase 1, chloroplastic | 3npe | The two parallel amphipathic helices (α1:85-109, α2:222-237) | L86, F87, R372, F439 | N85, L86, F87, Q88, A90, A91, A94, L95, A97, F98, G101, F102, V106, L107, A232, A235, C236, L371, R372 | N85, L86, F87, R89, A90, A91, A93, A94, L95, A97, F98, G101, F102, N105, V106, L107, P110, H221, A225, Y231, A232, A235, C236, I316, K317, L367, L371, R372, G373, G374, I438, F439, R559 | Bn [113, 114] |
| Monoglyceride lipase MGLL | 3jw8, chain A | Around amino acids L179, L186 | T168, F169, L172, L179, V180, L181 | T168, F169, K170, V171, L172, A173, A174, K175, V176, L177, N178, L179, V180, L181, P182, L184 | T168, F169, K170, L172, A173, K175, V176, L177, N178, L179, V180, L181, P182, N183, L184, S185, L186, G187 | Bn [115] |
| Dihydroorotate dehydrogenase | 3w7r | Region 31-68 | F37, H41, L49, W362 | F37, E40, H41, L42, P44, T45, L46, Q47, G48, L49, L50, L58, F62 | D34, F37, Y38, H41, L42, T45, L46, L49, L50, S54, R57, L58, R61, F62, L65, G66, L67, L68, R70, R245, R246, V247 | Fn [116, 117] |
| Phosphatase PTEN | 5bzz, chain A | Regions 260-269, 327-335 | L42, K263, M264 | V222, Q245, P248, T350, V351 | R41, L42, R47, R189, R335, V351 | Fn, Bn [118, 119] |
| (S)-mandelate dehydrogenase | 6bfg | Region 177-215 | Chain A: F6, R53, L54, Y179, L185, L195, H200 Chain B: W87, R172, K174, Y179, K182, H200 | Chain A: Y179, A181, V184, L185, C188, L189, P191, L195 Chain B: Y179, A181, V184, L185, C188, L189, P191, L195, V198, R199 | Chain A: Q3, N4, F6, R53, L54, D56, S178, Y179, S180, A181, V184, L185, G187, C188, L189, H190, P191, R192, S194, L195, F197, V198, R199, G201, Q358 Chain B: R172, S178, Y179, S180, A181, V184, | Bn [120] |

| | | | | | | |
|---|---|---|---|---|---|---|
| | | | | | L185, G187, C188, L189, P191, R192, S194, L195, V198, R199 | |
| (S)-mandelate dehydrogenase, homotetramer | Homotetramer biological assembly of 6bfg | Region 177-215 | Chain A: Y179, L195, H200<br>Chain B: Y179<br>Chain C: Y179, L195, H200<br>Chain D: Y179 | Chain A: Y179, A181, V184, L185, C188, L189, P191, L195, V198<br>Chain B: Y179, A181, V184, L185, C188, L189, P191, L195, V198, R199<br>Chain C: V184, L185, C188, L189, P191, L195<br>Chain D: V184, L185, C188 | Chain A:<br>S178, Y179, S180, A181, V184, L185, G187, C188, L189, H190, P191, R192, S194, L195, F197, V198, R199, G201<br>Chain B:<br>S178, Y179, S180, A181, V184, L185, G187, C188, L189, P191, R192, S194, L195, V198, R199<br>Chain C:<br>Y179, S180, A181, V184, L185, G187, C188, L189, P191, R192, S194, L195, V198, R199, G201<br>Chain D:<br>S178, Y179, S180, A181, V184, L185, G187, C188, P191, R192, S194, L195, V198, R199, G201 | Bn [120] |
| L-amino acid deaminase | 5hxw, chain A | I345, L347, L351, I352, F355 | F326, Y330, L333, L335, F355, M356 | V321, V322, K323, S325, F326, T327, Y330, L333, I345, S346, L347, N348, L351, I352, F355, M356 | V321, S325, F326, T327, G329, Y330, L333, P334, L336, A337, I345, L347, L351, I352, F355, M356 | Bn [121] |
| Intestinal fatty acid binding protein | 3akm, chain A | Around amino acid K27 | V26, K29 | V26, K29, L30, H33 | N24, I25,V26, K27, K29, L30 | Bn, FL [122, 123] |
| Phosphatidylinositol 4,5-bisphosphate 3-kinase | 4ovu | Kinase region 720-729, p110a region 863-872, and p85a region iSH2 | Chain A: M232, L233, W498, Y508, K867, G868, L870<br>Chain B: F512, M525 | Chain A: M232, L233<br>Chain B: - | Chain A: M232, L233, N331, R349, S379, N380, W498, S501, R502, E503, A504, G505, F506, Y508, K867, G868, A869<br>Chain B: F512, K519 | Fn [124] |
| Phosphatidylcholine transfer protein | 1ln1 | Region 184-193 | F107, W186, W190 | F107, P108 | P106, F107, P108, M109, S110, S147, G148, W186 | Fn [125] |

Bn – based on membrane binding affinities of mutants; FL – fluorescence; FLq – fluorescence quenching; Fn – functional studies of mutants; LD – linear dichroism measurements.

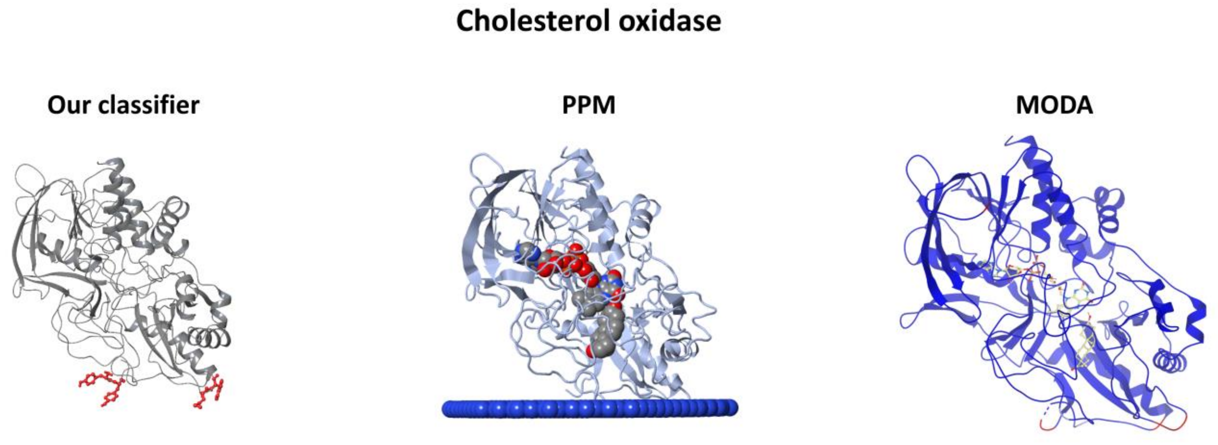


**Figure S11.** Comparison of the predictions provided from our classifier, PPM, and MODA for the cholesterol oxidase. Correct predictions for every tool.

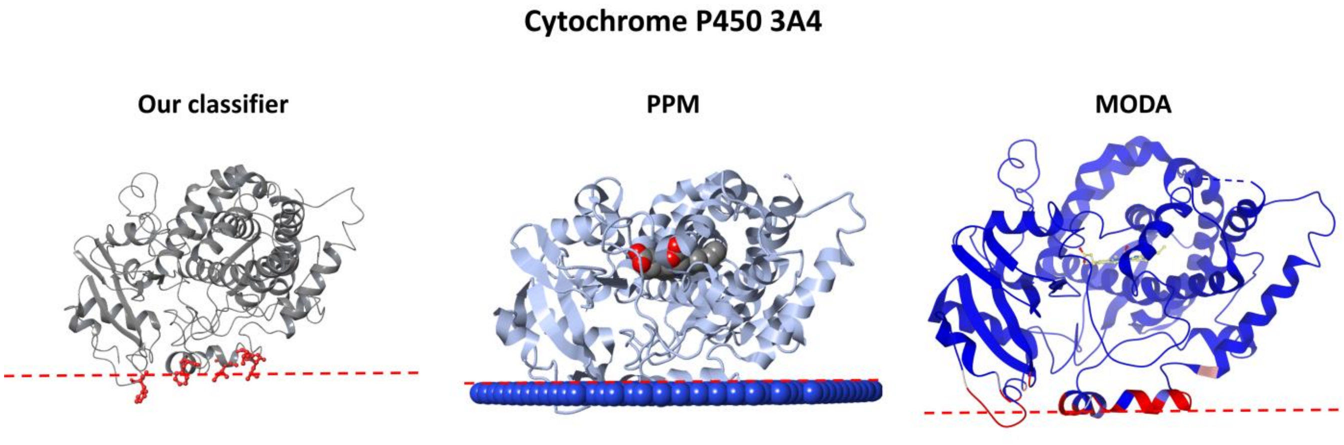


**Figure S12.** Comparison of the predictions provided from our classifier, PPM, and MODA for the cytochrome P450 3A4. Correct predictions for every tool. The membrane is depicted with a red dotted line according to Ref. [112].

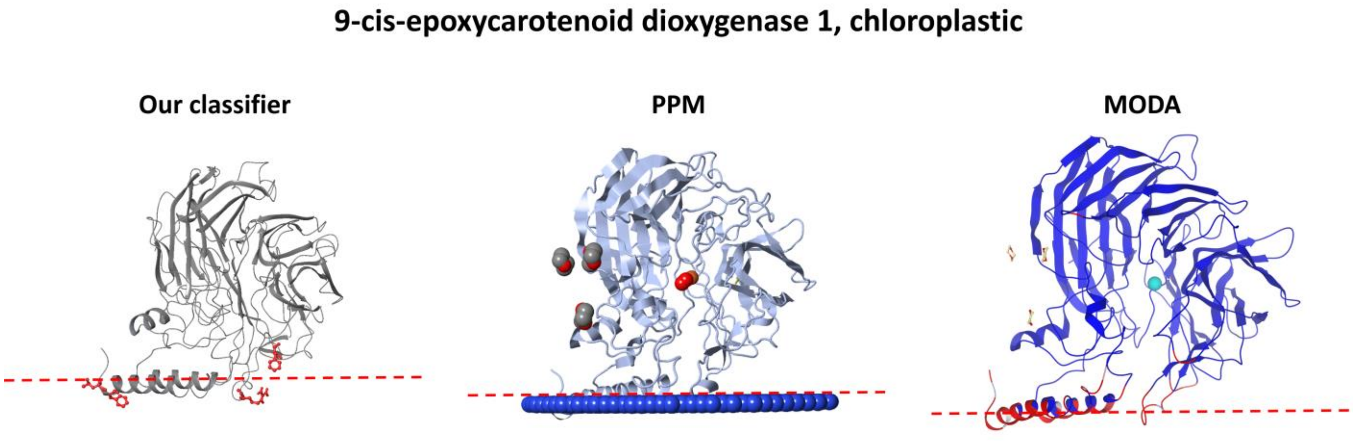


**Figure S13.** Comparison of the predictions provided from our classifier, PPM, and MODA for the 9-cis-epoxycarotenoid dioxygenase 1, chloroplastic. Correct predictions for every tool, with our classifier predicting the insertion of one of the two parallel amphipathic helices, instead of both. The membrane is depicted with a red dotted line according to Ref. [126].

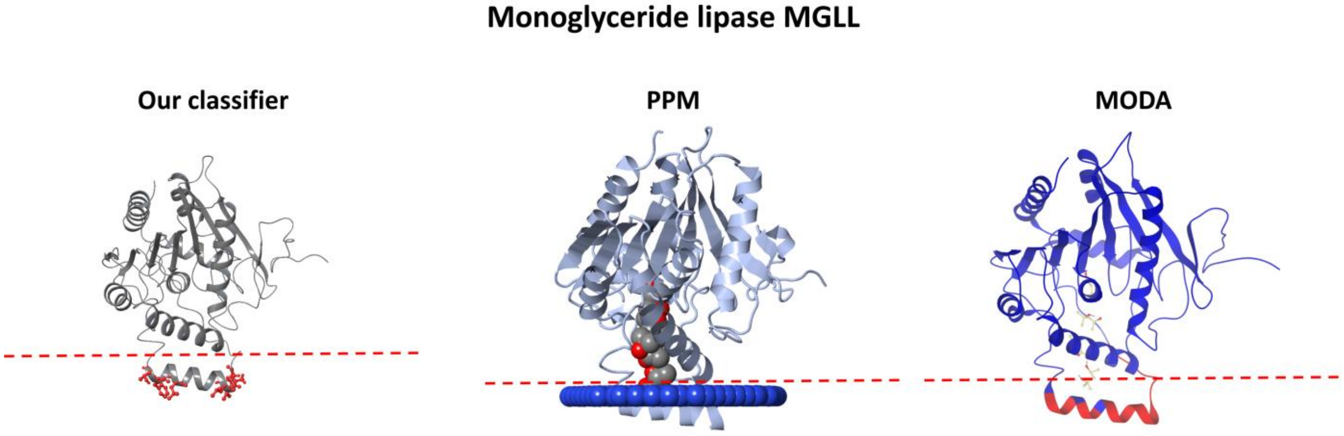


**Figure S14.** Comparison of the predictions provided from our classifier, PPM, and MODA for the monoglyceride lipase MGLL. Correct predictions for every tool. The membrane is depicted with a red dotted line according to Ref. [127].

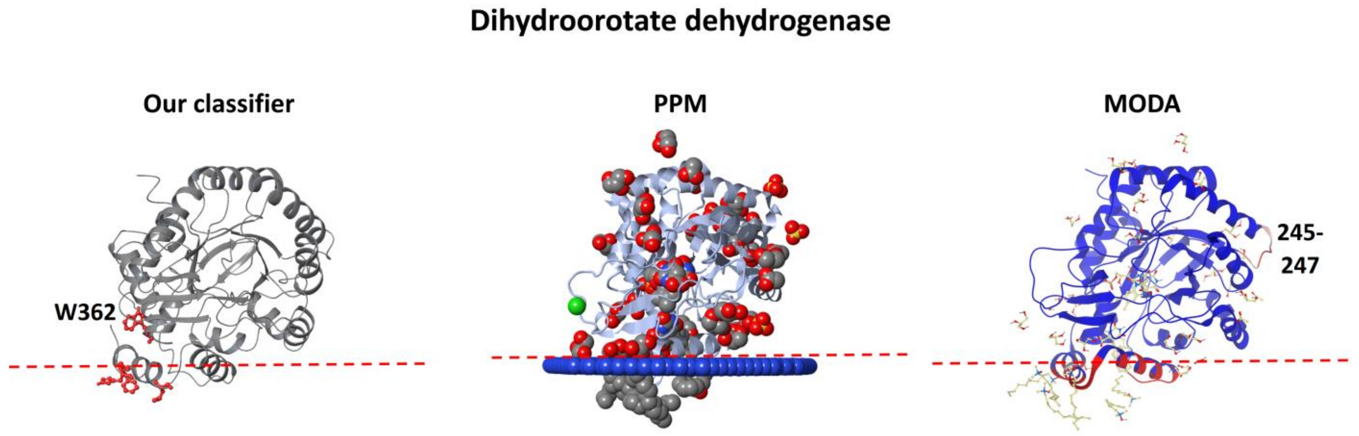


**Figure S15.** Comparison of the predictions provided from our classifier, PPM, and MODA for the dihydroorotate dehydrogenase. Correct predictions for every tool, with our classifier falsely identifying the amino acid W362 and MODA the region 245-247. The membrane is depicted with a red dotted line according to Ref. [117].

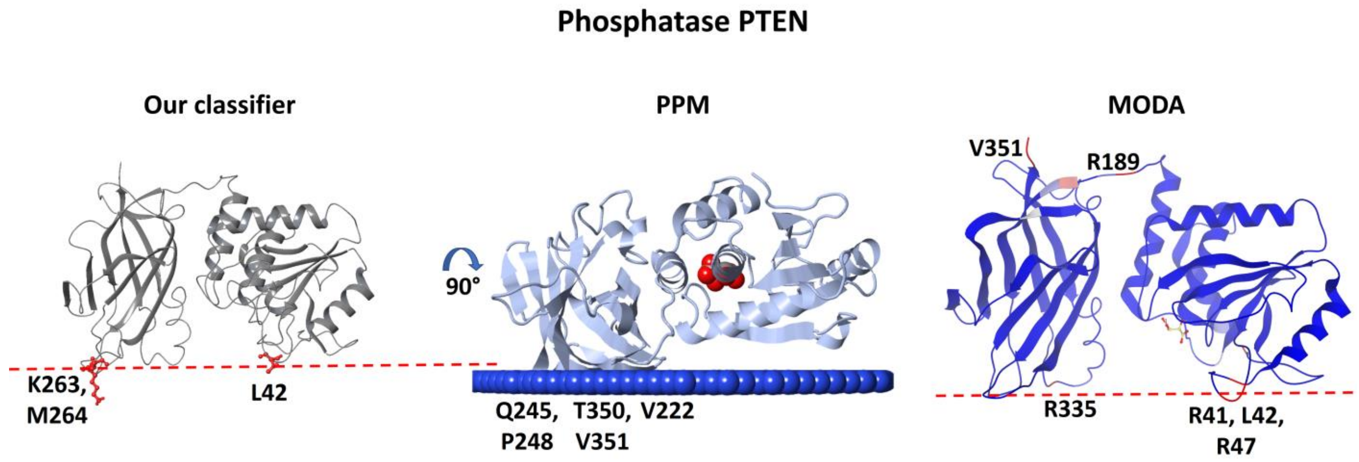


**Figure S16.** Comparison of the predictions provided from our classifier, PPM, and MODA for the phosphatase PTEN. Correct predictions for our classifier, with MODA correctly identifying the membrane binding region, but falsely identifying the opposite side of the C2 domain. PPM also falsely identified the opposite side of the C2 domain, providing a different orientation. The membrane is depicted with a red dotted line according to Ref. [119].

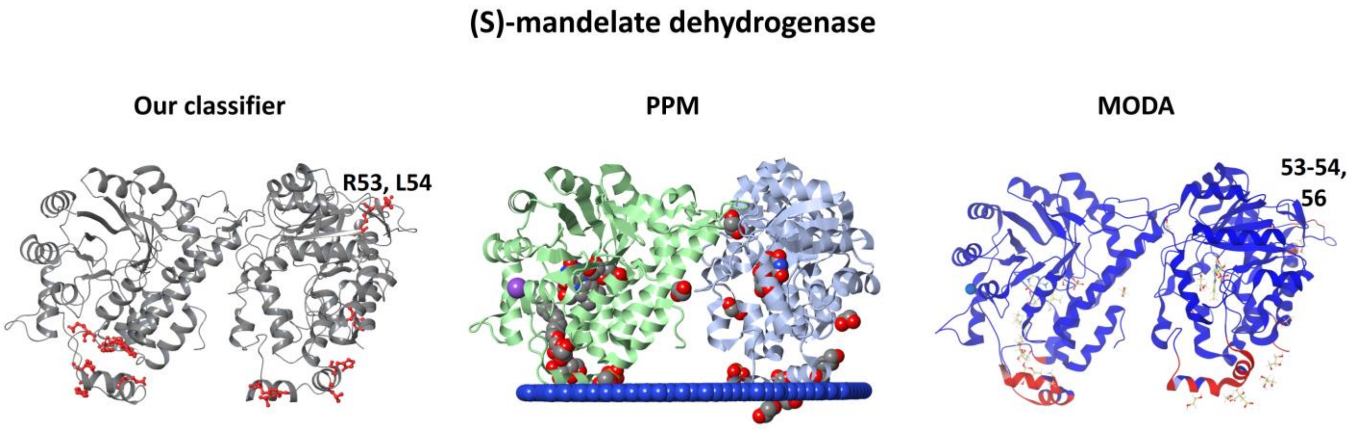


**Figure S17.** Comparison of the predictions provided from our classifier, PPM, and MODA for the (S)-mandelate dehydrogenase. Correct predictions for every tool, with our classifier and MODA falsely predicting the 53-56 region.

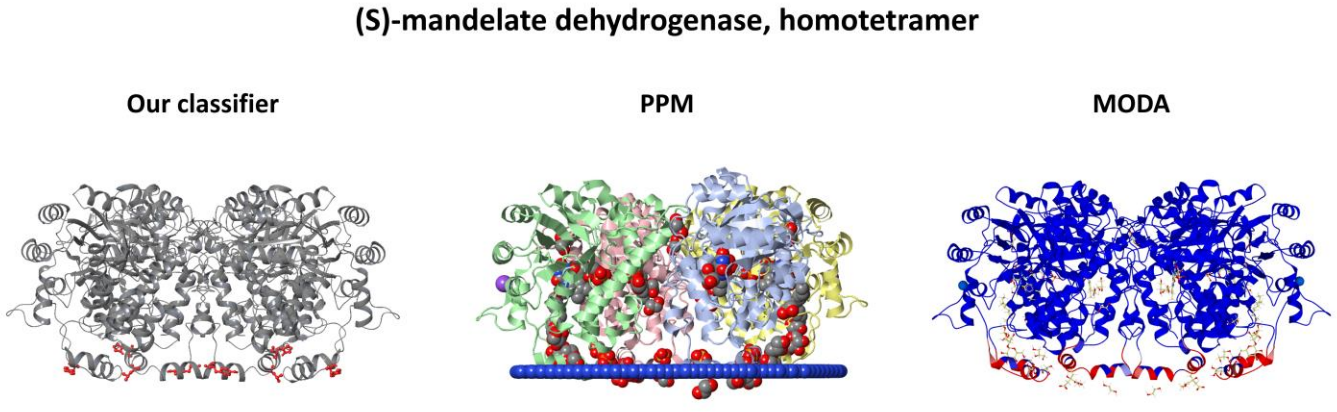


**Figure S18.** Comparison of the predictions provided from our classifier, PPM, and MODA for the homotetramer (S)-mandelate dehydrogenase (biological assembly). Correct predictions for every tool.

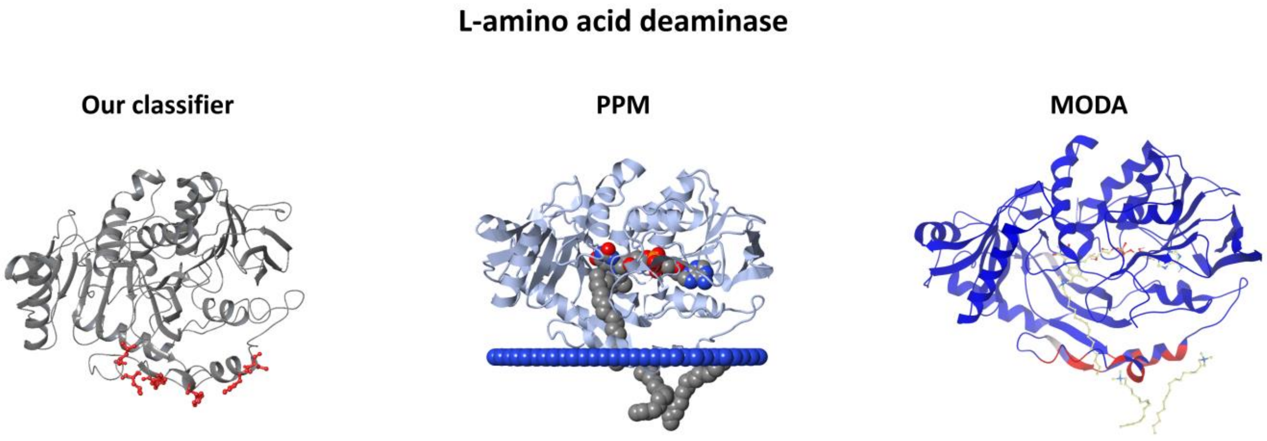


**Figure S19.** Comparison of the predictions provided from our classifier, PPM, and MODA for the L-amino acid deaminase. Correct predictions for every tool.

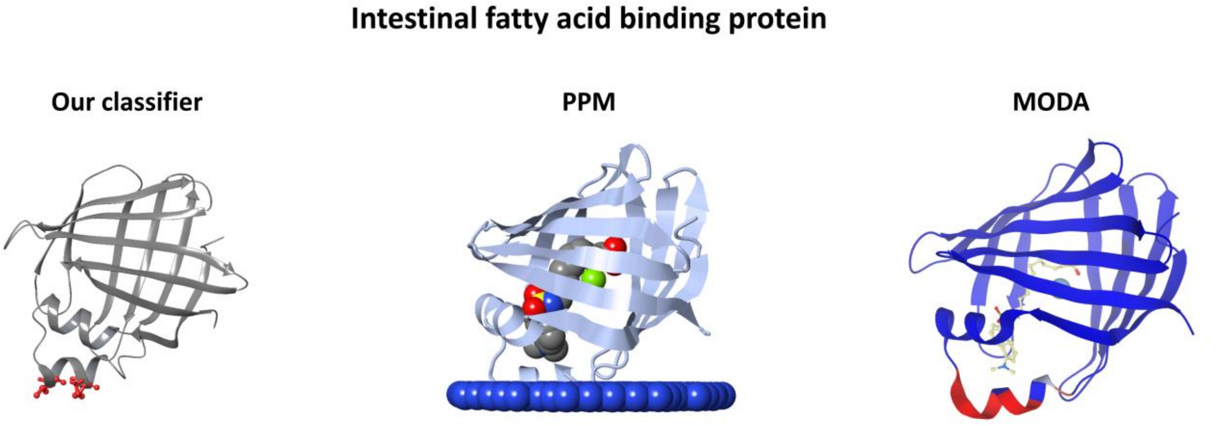


**Figure S20.** Comparison of the predictions provided from our classifier, PPM, and MODA for the intestinal fatty acid binding protein. Correct predictions for every tool.

### Phosphatidylinositol 4,5-bisphosphate 3-kinase

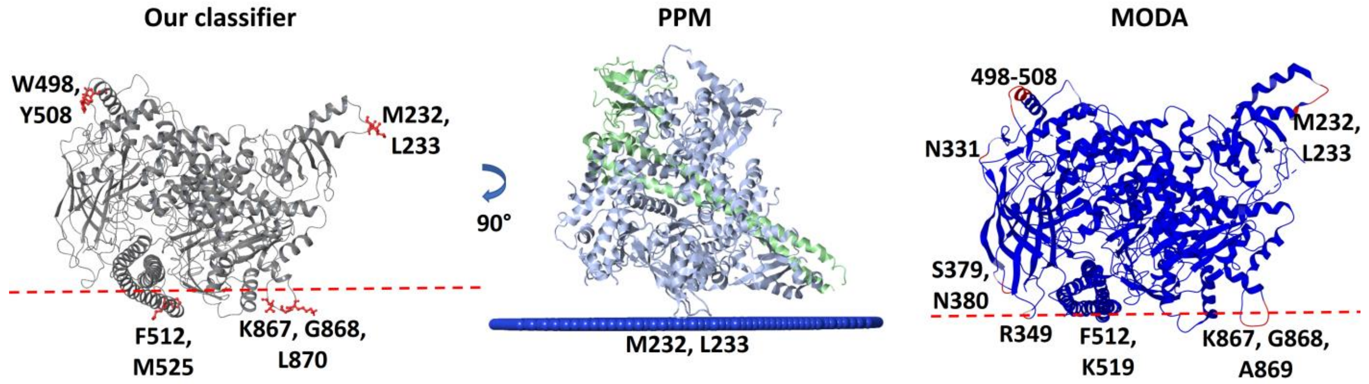


**Figure S21.** Comparison of the predictions provided from our classifier, PPM, and MODA for the phosphatidylinositol 4,5-bisphosphate 3-kinase. Every tool falsely predicted the region 232-233, with our classifier and MODA successfully identifying the p110a 863-872 region, but also having false positives the 498-508 region. PPM provides a membrane orientation different than the one proposed through mutagenesis experiments [124]. The membrane is depicted with a red dotted line according to Ref. [124].

### Phosphatidylcholine transfer protein

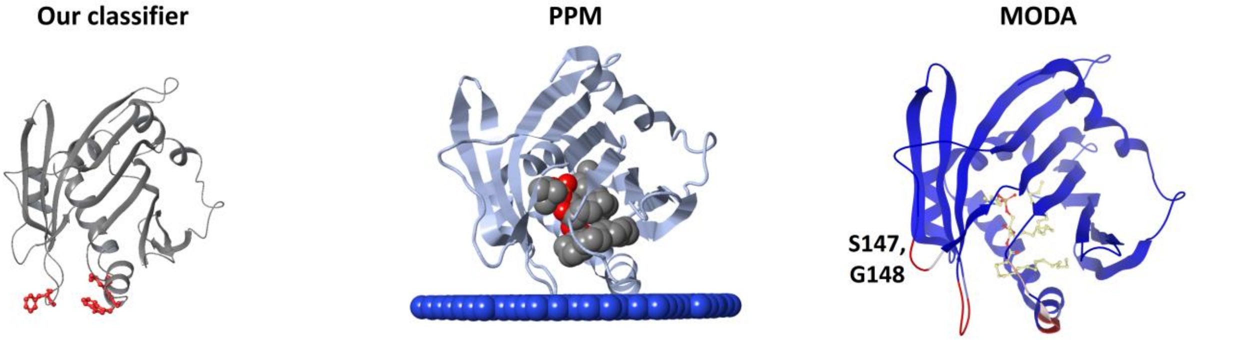


**Figure S22.** Comparison of the predictions provided from our classifier, PPM, and MODA for the phosphatidylcholine transfer protein. Correct predictions for every tool, with MODA suggesting also the insertion of the 147-148 loop.

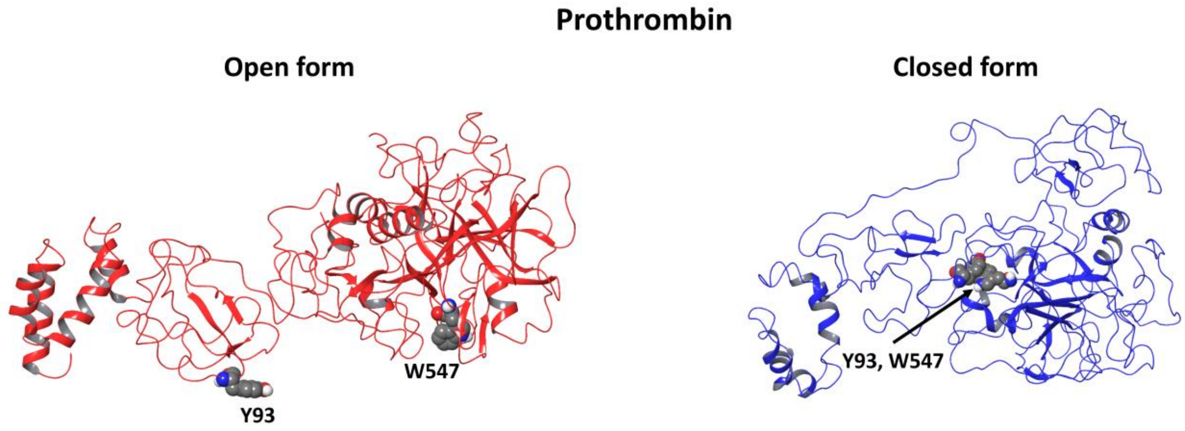


**Figure S23.** The open (PDB: 5EDM [128]) and closed (PDB: 6BJR [129]) forms of the prothrombin protein. The amino acids Y93 and W547 (W533 of 5EDM) are highlighted in CPK representation. In the closed form, Y93 inserts into the active site of the protease domain where it interacts with W547, while in the open form is distant from W547, suggested from our classifier and MODA to insert in the membrane.

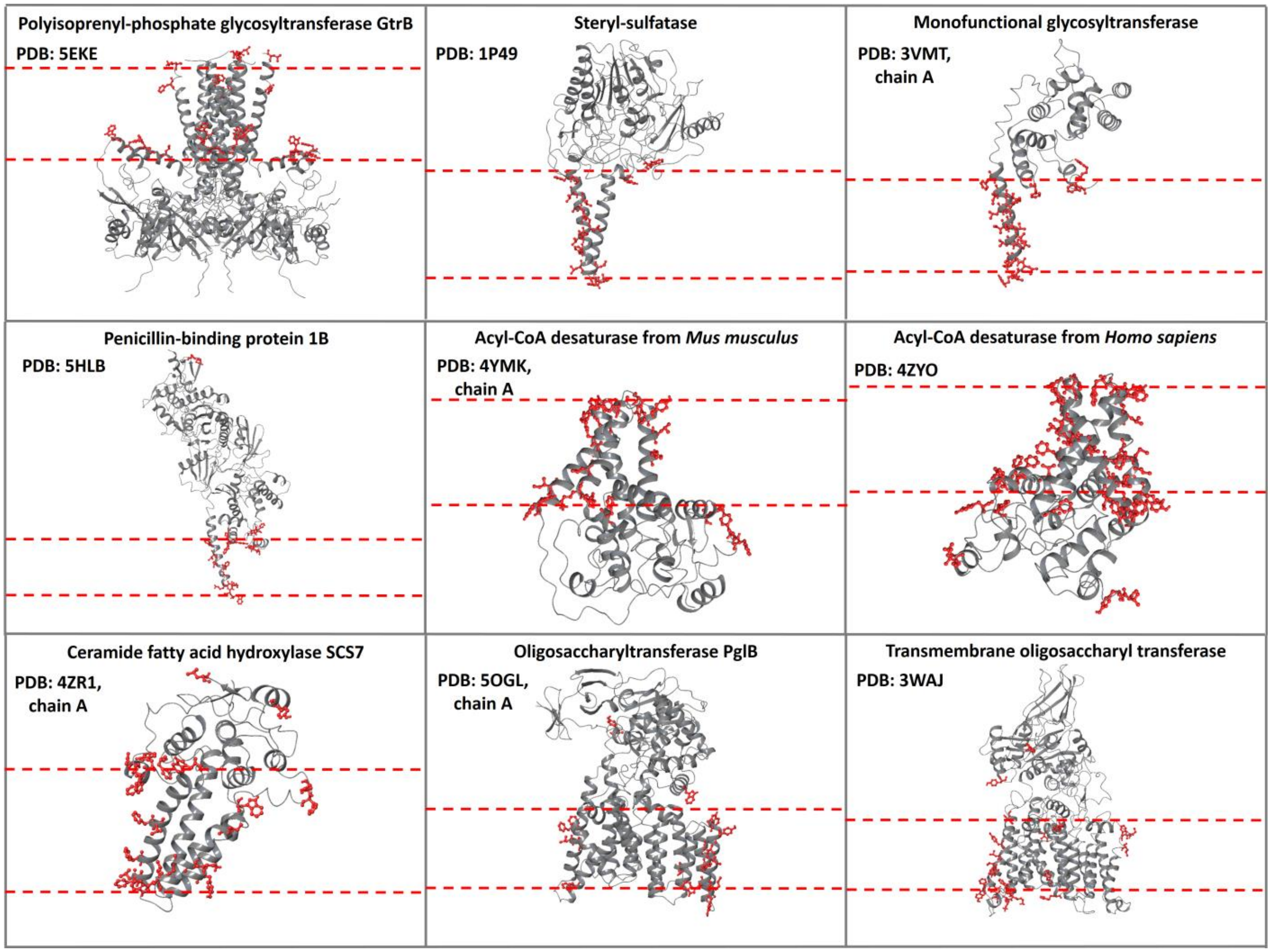


**Figure S24.** Our classifier's predictions in nine transmembrane enzymes with soluble domain performing extracellular catalysis. The membrane-penetrating amino acids predicted from our classifier are depicted with red, and the membrane is depicted with red dotted lines. The proteins are placed in the membrane according to available structural information [130]. Overall, the predictions are in agreement with the experiments, except for some use cases where a few predictions are false positives.
